# A comprehensive assessment of inbreeding and laboratory adaptation in *Aedes aegypti* mosquitoes

**DOI:** 10.1101/237776

**Authors:** Perran A. Ross, Nancy M. Endersby-Harshman, Ary A. Hoffmann

**Author notes:** Email addresses: Perran A. Ross, Nancy M. Endersby-Harshman, Ary A. Hoffmann.

## Abstract

Modified *Aedes aegypti* mosquitoes reared in laboratories are being released around the world to control wild mosquito populations and the diseases they transmit. Several efforts have failed due to poor competitiveness of the released mosquitoes. We hypothesized that colonized mosquito populations could suffer from inbreeding depression and adapt to laboratory conditions, reducing their performance in the field. We established replicate populations of *Ae. aegypti* mosquitoes collected from Queensland, Australia, and maintained them in the laboratory for twelve generations at different census sizes. Mosquito colonies maintained at small census sizes (≤100 individuals) suffered from inbreeding depression due to low effective population sizes which were only 25% of the census size as estimated by SNP markers. Populations that underwent full-sib mating for 9 consecutive generations had greatly reduced performance across all traits measured. We compared the established laboratory populations with their ancestral population resurrected from quiescent eggs for evidence of laboratory adaptation. The overall performance of laboratory populations maintained at a large census size (400 individuals) increased, potentially reflecting adaptation to artificial rearing conditions. However most individual traits were unaffected, and patterns of adaptation were not consistent across populations. Differences between replicate populations may indicate that founder effects and drift affect experimental outcomes. Though we find limited evidence of laboratory adaptation, mosquitoes maintained at low population sizes can clearly suffer fitness costs, compromising the success of “rear and release” strategies for arbovirus control.

## Introduction

*Aedes aegypti* mosquitoes transmit some of the most important arboviruses in the world, including dengue, Zika and chikungunya. These diseases are an enormous burden to global health and the eradication or disruption of their vectors is currently the leading approach to their control. Several of these strategies rely on rearing and releasing modified mosquitoes into the environment to reduce disease incidence. The sterile insect technique has been used for decades to suppress mosquito populations, though many programs using this technique have not succeeded in achieving substantial population suppression [1–3]. In this approach, male mosquitoes are irradiated or chemically treated and then released into the field in large numbers to sterilize the wild females. Alternatives to this technique have recently emerged which do not rely on traditional sterilization (reviewed in [4, 5]). Transgenic *Ae. aegypti* males possessing a dominant lethal system have been released in multiple locations where they have reduced population sizes, at least in the short term [6–9]. When these males mate with wild females, most offspring die before reaching the late pupal stage, though a low proportion can emerge as functional adults [10] and may persist for months after releases cease [9]. *Aedes* mosquitoes infected experimentally with the intracellular bacterium *Wolbachia* are also being released into the field for disease control programs. Certain strains of *Wolbachia* reduce the capacity for mosquitoes to transmit RNA viruses [11] and infected males can sterilize wild, uninfected females through cytoplasmic incompatibility [12, 13]. Mosquitoes infected with *Wolbachia* are now being released into the field, both to suppress populations [14, 15] and to replace populations with mosquitoes that are refractory to virus transmission [16–18].

Rear and release approaches to arbovirus control require large quantities of mosquitoes to be reared in the laboratory for eventual release into the field. For sterile and incompatible male approaches, high ratios of modified to wild males are needed to achieve suppression, particularly if the modifications have deleterious effects on male fitness [19]. Laboratory environments are inherently artificial, and colonized mosquito populations experience an entirely different set of selective pressures compared to natural populations [20]. Many laboratory mosquito populations are held at a controlled temperature, humidity and photoperiod, provided with abundant nutrition, and reared in discrete generations according to a schedule (e.g. [21–23]). Rearing insects in discrete generations may select for an earlier, shorter and more productive reproductive period, as only individuals that adhere to the rearing schedule will contribute to the next generation [24, 25]. Laboratory populations of insects are often maintained at high adult densities due to space limitation which could lead to intense male-male competition and altered courtship behaviour [26–28]. Laboratory environments can also lack selective pressures which could lead to declines in later life reproduction [29], a reduced ability to survive temperature extremes, dry conditions or starvation [30] or a loss of insecticide resistance [31]. Maintaining populations in the laboratory can also cause a reduction in genetic diversity resulting in low adaptive potential and inbreeding depression [32]. Laboratory environments can therefore impose rapid genetic changes on insect populations, and laboratory-derived mosquitoes could be mal-adapted to the target population when eventually released into the field [33].

Competitive mosquitoes are critical for the success of rear and release programs. Past sterile insect interventions have failed due to the poor performance of released mosquitoes, possibly caused by laboratory adaptation [1, 34, 35]. For releases of sterile, dominant-lethal or incompatible males, the ability of modified males to seek and inseminate wild females is especially important [36, 37]. For approaches where modified mosquitoes are intended to persist in the environment, it is often necessary for them to perform similarly to wild mosquitoes. Two attempts to establish the *w*MelPop *Wolbachia* infection in natural *Ae. aegypti* populations failed due to deleterious effects associated with the infection, including costs to fecundity, adult lifespan and egg viability [17]. While trait variation related to fitness in mosquitoes has often been well-characterized, there are fewer attempts to compare laboratory strains intended for release against the wild mosquitoes against which they are intended to compete.

Across all mosquito species, there are numerous studies that compare life history, morphological and physiological traits between laboratory and field populations for evidence of laboratory adaptation (S1 Table). Substantial and rapid adaptation by mosquitoes to laboratory conditions is often observed (e.g. [38, 39]), but there are several instances of laboratory populations suffering reduced fitness (e.g. [40, 41]). Other studies find no clear differences between laboratory and field populations despite years of separation (e.g. [42–44]). Mosquitoes maintained in the laboratory can differ from wild populations for many traits, including blood-feeding duration [45, 46], wing shape [47], oviposition behaviour [48], mating success [49–51], swarming behaviour [26] and susceptibility to pathogens [52–55]. Researchers often compare a single wild population to a single long-established laboratory population (e.g. [49, 51]), but these results could be confounded by inbreeding, drift and bottlenecks in the laboratory population rather than reflecting laboratory adaptation. Differences between populations could also be affected by rearing conditions, for example if the wild population is reared under field conditions (e.g. [39, 40, 56] or if measurements are conducted at different time points (e.g. [46, 52]). Other studies compare populations collected from different locations (e.g. [48, 53, 57]) and any effects of laboratory maintenance could be confounded by local adaptation.

The extent of laboratory adaptation can vary between insect orders (Hoffmann and Ross unpublished) and this could reflect differences in the range of conditions that can be tolerated relative to the conditions experienced in the laboratory [58]. Laboratory environments that are suboptimal will impose strong selective pressures on mosquito populations, leading to rapid adaptation (e.g. [38]). Colonized mosquito species can require a specific set of conditions such as swarm markers [38] artificial horizons [59], dusk periods [59] or exposure to stroboscopic light [60] to improve their reproductive success in the laboratory. Other species will not freely reproduce in the laboratory at all, requiring induced copulation over successive generations before free-mating colonies can be established [61, 62]. In contrast, *Ae. aegypti* collected from the field perform well in the laboratory without any of these specific requirements (e.g. [22]), and therefore less drastic differences in traits would be expected between laboratory and field populations due to a lack of selective pressures.

Rear and release programs with modified *Ae. aegypti* mosquitoes are now underway in several countries, and many of these programs rely on the use of mosquitoes that have been inbred or maintained in the laboratory for extended periods. We colonized replicate *Ae. aegypti* populations collected from Queensland, Australia to assess the effects of laboratory maintenance and inbreeding on life history traits in this species. We find that inbreeding is costly and is associated with a reduction in effective population size, but we find limited evidence of laboratory adaptation for most life history traits. Modified mosquitoes reared for disease control programs should therefore be maintained at large population sizes and/or crossed with field populations prior to field release. Our research highlights potential issues with maintaining colonized insects that are destined for field release, and inform protocols for the maintenance of *Ae. aegypti* in the laboratory.

## Results

### Population establishment and preliminary fitness comparisons

We established a population of *Ae. aegypti* in the laboratory with eggs collected from Townsville, Australia in 2015. Mosquitoes were maintained using a protocol designed to minimize selective pressures [63]. After one generation in the laboratory, we split the population into replicate cages that were maintained at different census sizes (Figure 1, see *Materials and methods*). Five populations were maintained at a census size of 400 adults (large populations) and five populations were maintained at a census size of 100 adults (small populations). We also established five isofemale lines and ten lines which were maintained with a single male and female each generation (inbred lines). At F_2_, we compared a range of life history traits with an established laboratory population from Cairns (F_11_), however we observed no differences between populations for any trait (S1 Appendix).

**Figure 1.**
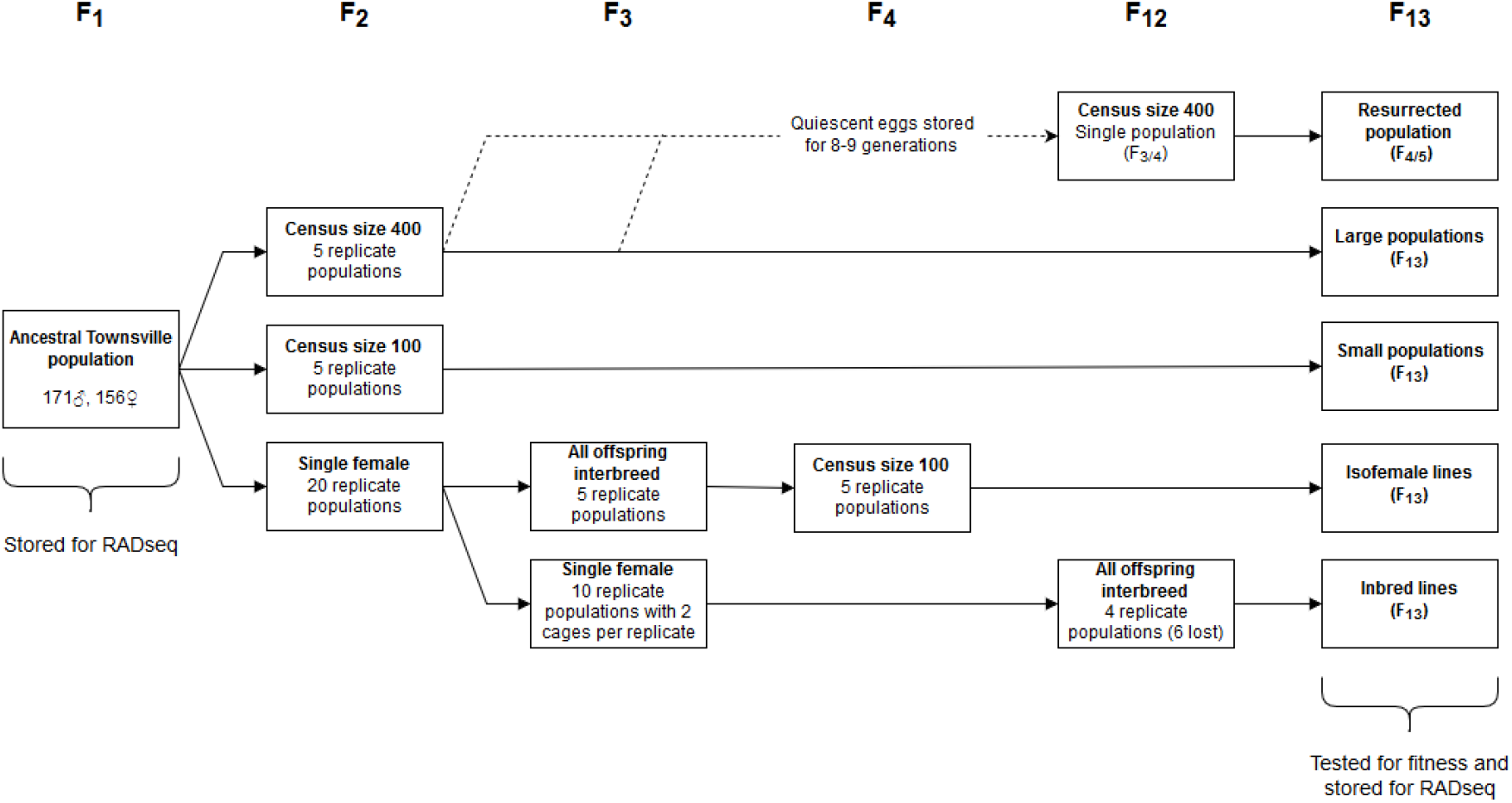
Maintenance scheme for replicate *Ae. aegypti* laboratory populations. An ancestral population was established from eggs collected from Townsville, Australia that all other populations were derived from. Replicate populations were maintained separately beginning from F_2_ and were not interbred.

At F_5_, we compared life history traits between replicated large populations and inbred lines. After two generations of brother-sister mating, the inbred lines had reduced fitness relative to the large populations, with substantial costs to larval survival and development time (S2 Appendix). We also observed significant differences between replicate populations for some life history traits which likely arose due to founder effects or drift (S2 Appendix).

### Fitness comparisons at F_13_

When the Townsville populations had reached F_13_, we performed fitness comparisons to test for inbreeding effects, laboratory adaptation, drift and founder effects. The original Townsville population was resurrected from quiescent eggs to enable concurrent comparisons between the laboratory populations at F_13_ and their ancestor at F_4/5_ to test for laboratory adaptation. Populations from Cairns at F_2_ and F_22_, and populations from Innisfail at F_4_ and F_10_ were also included in all comparisons. Not all inbred lines were included in the experiments as the majority were lost by F_13_ (S2 Table).

We measured larval development time for all populations under high nutrition and low nutrition conditions (Figure 2). Under high nutrition conditions, small populations were developmentally delayed compared to large populations (females: one-way ANOVA: F_1,38_ = 48.080, P < 0.001, males: F_1,38_ = 27.895, P < 0.001). There were significant differences in development time between replicate cages of the large populations (females: F_4,15_ = 4.262, P = 0.017, males: F_4,15_ = 4.699, P = 0.012), isofemale lines (females: F_4,15_ = 81.888, P < 0.001, males: F_4,15_ = 18.956, P < 0.001), inbred lines (females: F_3,9_ = 10.413, P = 0.003, males: F_3,9_ = 8.575, P = 0.005) and also females from the small populations (F_4,15_ = 6.749, P = 0.003) but not males (F_4,15_ = 2.031, P = 0.141). The isofemale and inbred lines were particularly diverse; some lines performed as well as (or better than) the large populations while others had greatly extended development times (Figure 2). Large populations (Townsville F_13_) were consistently faster to develop than the Townsville F_4/5_ ancestral population (females: F_1,22_ = 33.462, P < 0.001, males: F_1,22_ = 15.434, P = 0.001), suggesting a positive effect of laboratory maintenance on this trait. The Cairns F_22_ laboratory population was also faster to develop than the Cairns F_2_ field population (females: F_1,6_ = 6.407, P = 0.045, males: F_1,6_ = 9.147, P = 0.023), though the opposite was true for Innisfail, where the laboratory population was slower to develop. This difference was however only significant for females (females: F_1,6_ = 6.653, P = 0.042, males: F_1,6_ = 5.938, P = 0.051). Under low nutrition conditions, development times were greatly extended relative to high nutrition conditions (Mann-Whitney U: Z = 16.250, P < 0.001, Figure 2). Differences between groups of populations maintained at different census sizes became less clear as there were significant differences between replicate populations at all census sizes (one-way ANOVA: all P ≤ 0.002).

**Figure 2.**
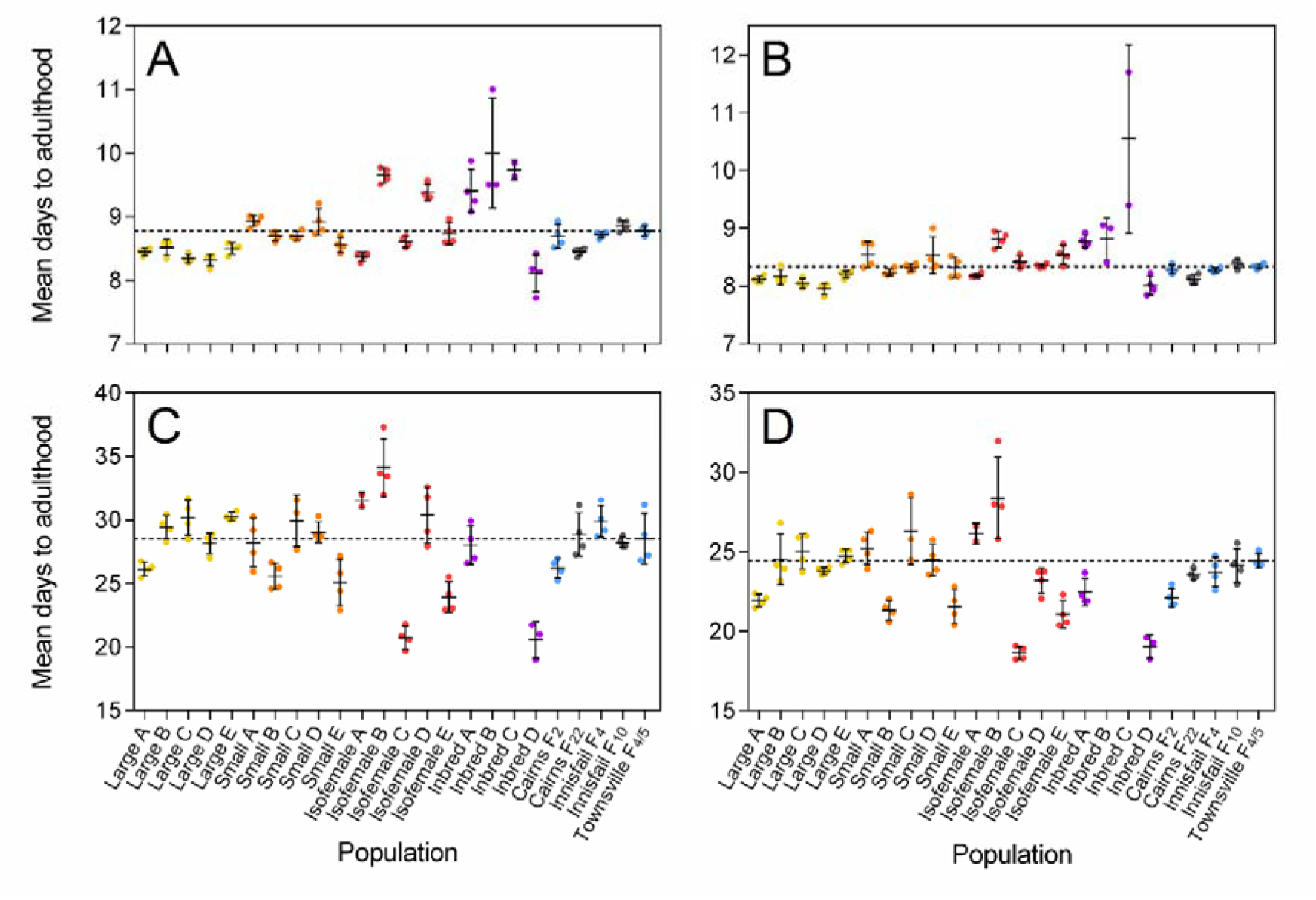
Development time of *Aedes aegypti* F_13_ laboratory populations maintained at different census sizes. Mean development time was measured for (A&C) female and (B&D) male larvae under (A&B) high nutrition (food provided *ad libitum)* and (C&D) low nutrition (0.1 mg of TetraMin^®^ per larva every 2 days) conditions. Each data point represents the mean development time of individuals from a single container of 100 larvae. Replicates of the large populations (Townsville F_13_, census size 400) are in yellow, small populations (Townsville F_13_, census size 100) in orange, isofemale lines (Townsville F_13_) in red and inbred lines (Townsville F_13_) in purple. Other laboratory populations are shown in gray and ancestral / field populations in pale blue. Inbred lines B and C were not tested under low nutrition conditions. The dashed line represents the mean development time of the Townsville F_4/5_ ancestral population. Error bars are standard deviations.

We compared the proportion of larvae that survived to adulthood between populations in the larval development experiment (Figure 3). Under high nutrition conditions, survival rates approached 100% in most populations that were maintained at a census size of 400. Small populations had reduced survival rates compared to large populations (Mann-Whitney U: Z = 3.273, P = 0.001), as did the isofemale (Z = 3.489, P < 0.001) and inbred (Z = 4.018, P < 0.001) lines. We observed significant variation between isofemale lines (Kruskal-Wallis: χ^2^ = 14.872, df = 4, P = 0.005), but not between large (χ^2^ = 2.129, df = 4, P = 0.712) or small (χ^2^ = 7.507, df = 4, P = 0.111) populations. Survival to adulthood was poorer under low nutrition conditions (Mann-Whitney U: Z = 6.343, P < 0.001), but populations maintained at a census size of 400 still performed consistently better than populations maintained at lower census sizes (Z = 7.084, P < 0.001). No differences between laboratory and field populations from Townsville, Innisfail or Cairns were evident (Mann-Whitney U: all P > 0.05), but the ability to detect any differences with this test was low due to low sample sizes for each population. Sex ratios of adults emerging from the larval development experiment did not deviate significantly from 1:1 under high nutrition conditions for all populations (Chi-square: df = 3, all P > 0.05), except for the Cairns F_2_ population which was biased towards males (df = 3, P = 0.013). Sex ratios were skewed towards males under low nutrition conditions (df = 86, P < 0.001) which was likely the result of female larval mortality.

**Figure 3.**
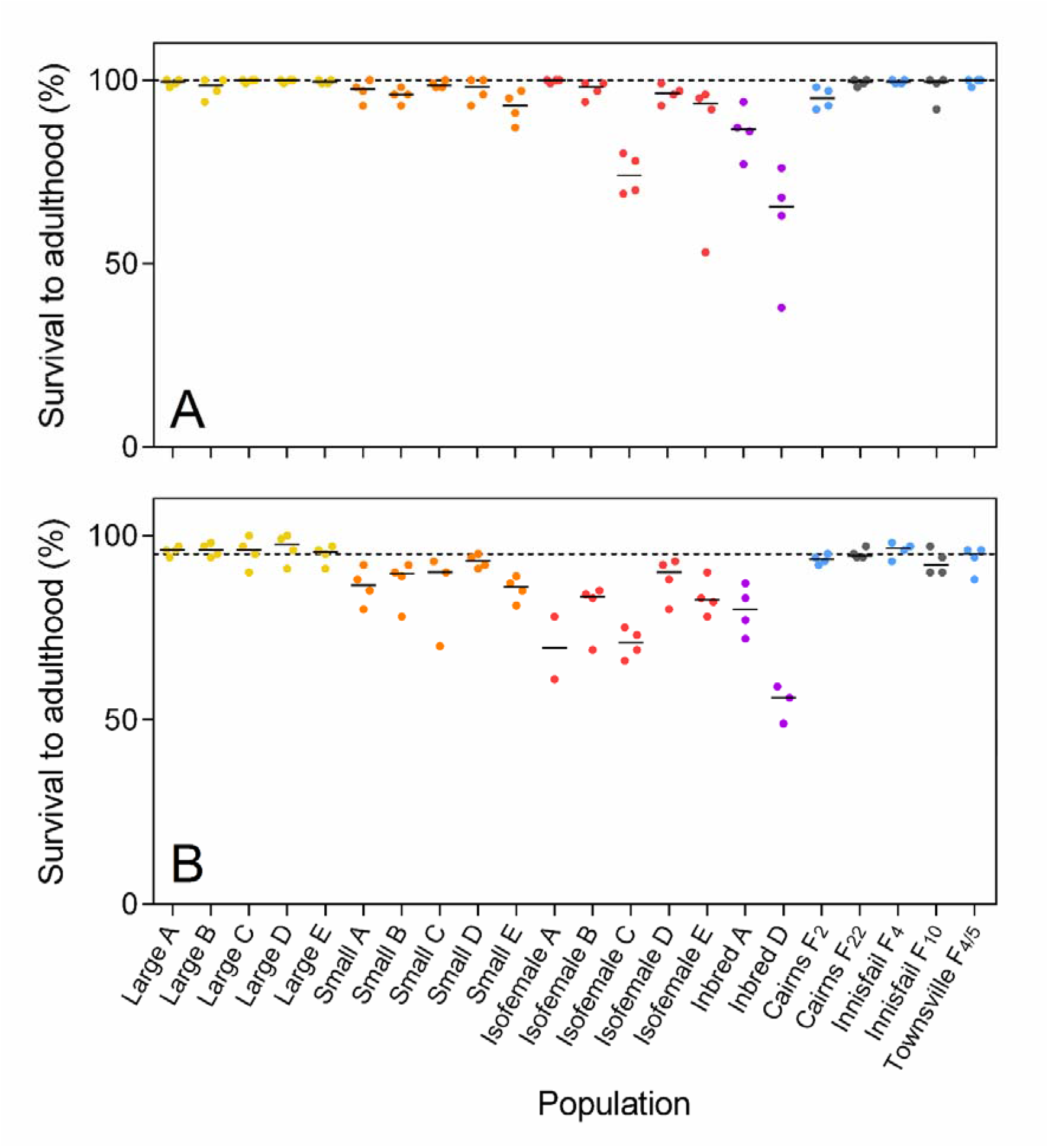
Survival to adulthood of *Aedes aegypti* F_13_ laboratory populations maintained at different census sizes. The percentage of larvae surviving to adulthood was tested under (A) high nutrition (food provided *ad libitum*) and (B) low nutrition (0.1 mg of TetraMin^®^ per larva every 2 days) conditions. Replicates of the large populations (Townsville F_13_, census size 400) are in yellow, small populations (Townsville F_13_, census size 100) in orange, isofemale lines (Townsville F_13_) in red and inbred lines (Townsville F_13_) in purple. Other laboratory populations are shown in gray and ancestral / field populations in pale blue. Solid black lines indicate the median survival of each population. The dashed line represents the median survival of the Townsville F_4/5_ ancestral population.

A random subset of females emerging from the larval development experiment was scored for their fecundity (Figure 4) and egg hatch proportion (Figure 5). Patterns of fecundity were similar between the first and second gonotrophic cycles, but fecundity was lower overall in the second gonotrophic cycle (one-way ANOVA: F_1,624_ = 17.660, P < 0.001). We considered only the first gonotrophic cycle for the following analyses, as some females died before the second gonotrophic cycle. Inbred populations had greatly reduced fecundity compared to large populations (F_1,100_ = 130.395, P < 0.001). Replicate populations differed significantly from each other for large populations (F_4,69_ = 3.573, P = 0.010), isofemale lines (F_4,68_ = 10.300, P < 0.001) and inbred lines (F_3,24_ = 12.087, P < 0.001), but not small populations (F_4,66_ = 1.677, P = 0.166), potentially reflecting drift or founder effects. The fecundity of large populations (Townsville F_13_) did not differ from that of Townsville F_4/5_ (F_1,86_ = 0.549, P = 0.461), indicating that the effects of laboratory maintenance on this trait are minimal.

**Figure 4.**
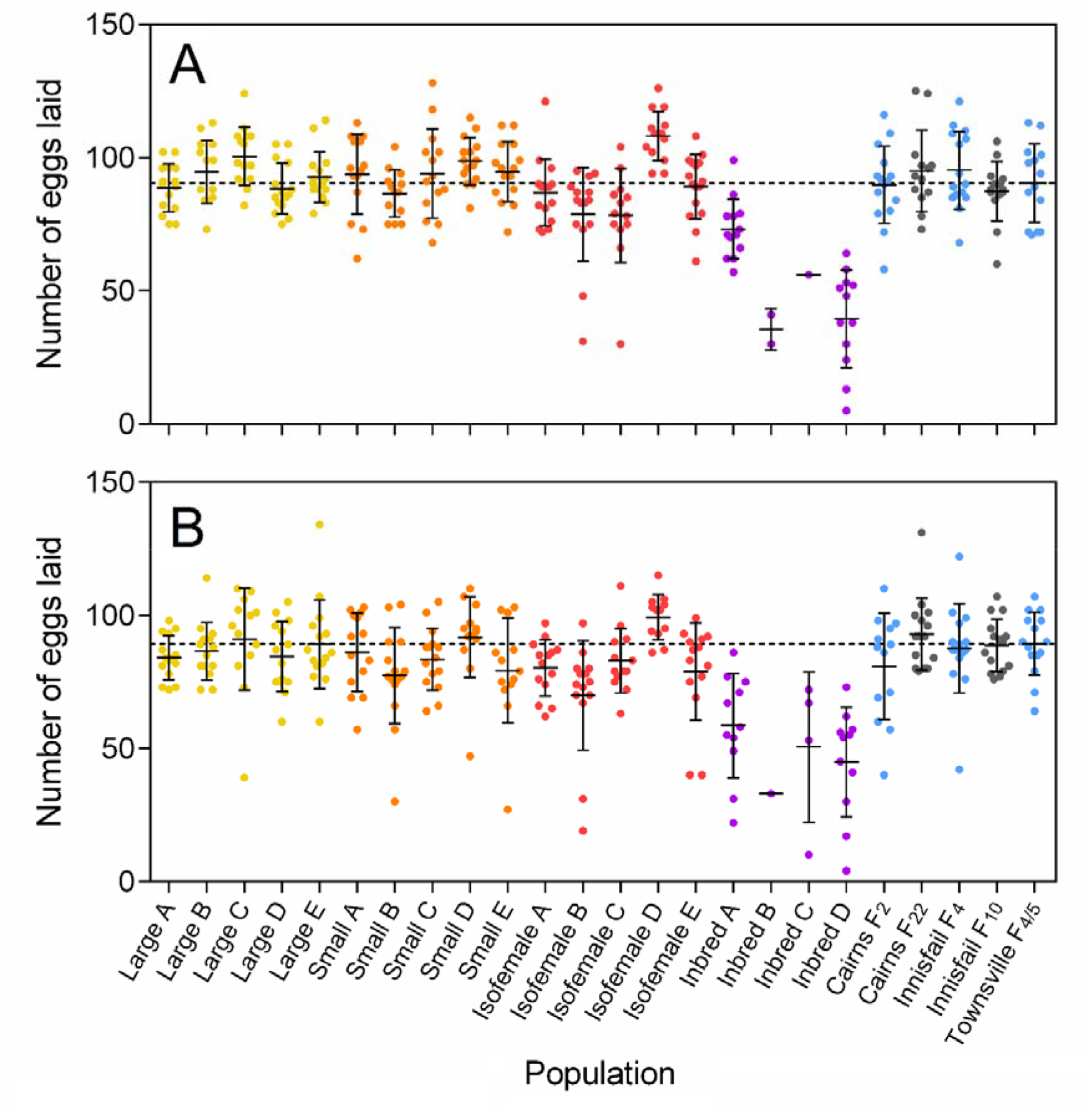
Fecundity of *Aedes aegypti* F_13_ laboratory populations maintained at different census sizes. The number of eggs laid by females in each experimental population was scored during their first (A) and second (B) gonotrophic cycle. Fifteen females were tested per line, or as many as possible for inbred lines B and C. Replicates of the large populations (Townsville F_13_, census size 400) are in yellow, small populations (Townsville F_13_, census size 100) in orange, isofemale lines (Townsville F_13_) in red and inbred lines (Townsville F_13_) in purple. Other laboratory populations are shown in gray and ancestral / field populations in pale blue. The dashed line represents the mean fecundity of the Townsville F_4/5_ ancestral population. Error bars are standard deviations.

**Figure 5.**
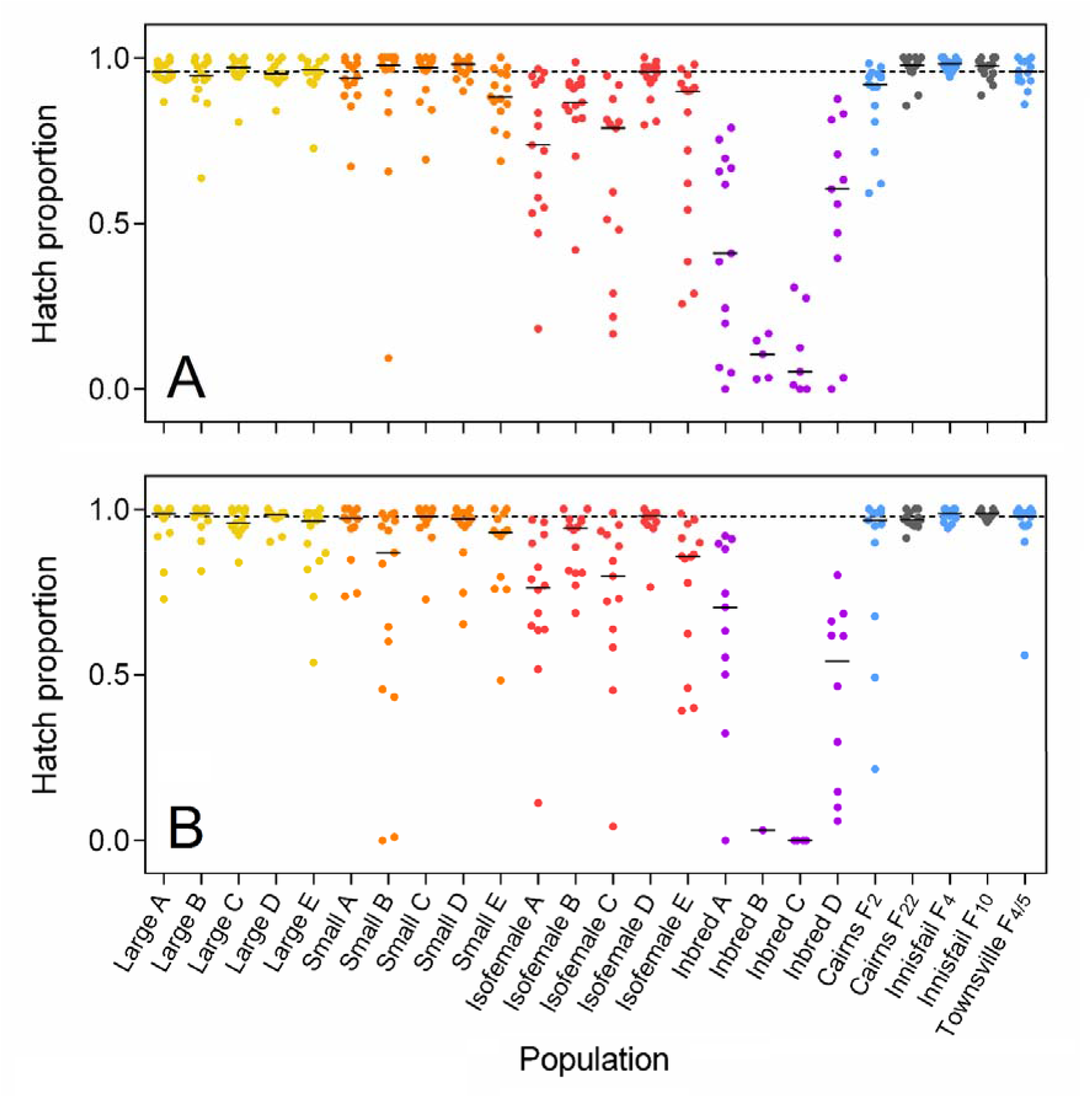
Egg hatch proportions of *Aedes aegypti* F_13_ laboratory populations maintained at different census sizes. Egg hatch proportions in each experimental population were scored in their first (A) and second (B) gonotrophic cycle. Fifteen females were tested per line, or as many as possible for inbred lines B and C. Replicates of the large populations (Townsville F_13_, census size 400) are in yellow, small populations (Townsville F_13_, census size 100) in orange, isofemale lines (Townsville F_13_) in red and inbred lines (Townsville F_13_) in purple. Other laboratory populations are shown in gray and ancestral / field populations in pale blue. Solid black lines indicate the median egg hatch proportion of each population. The dashed line represents the median egg hatch proportion of the Townsville F_4/5_ ancestral population.

Egg hatch proportions did not differ between gonotrophic cycles (Mann-Whitney U: Z = 1.773, P = 0.077), but were substantially affected by inbreeding, with both isofemale lines (Z = 6.895, P < 0.001) and inbred lines (Z = 8.334, P < 0.001) exhibiting reduced hatch proportions relative to large populations (Figure 5). There were differences between replicate populations for small populations (Kruskal-Wallis: χ^2^ = 10.405, df = 4, P = 0.034), isofemale (χ^2^ = 19.639, df = 4, P = 0.001) and inbred lines (χ^2^ = 11.222, df = 3, P = 0.011), but not large populations (χ^2^ = 3.141, df = 4, P = 0.535). Hatch proportions did not differ between the Townsville F_4/5_ population and the large populations at F_13_ (Mann-Whitney U: Z = 0.2137, P = 0.834), but were improved in the Cairns F22 population relative to Cairns F_2_ (Z = 3.202, P = 0.001), suggesting a positive effect of laboratory maintenance.

We measured wing length from a random subset of adults emerging from the larval development experiment to estimate the body size of each population under different nutritional conditions (Figure 6). Wing lengths under high nutrition conditions were much larger than under low nutrition conditions (one-way ANOVA: F_1,918_ = 221.749, P < 0.001) and differences between populations were more distinct. Under high nutrition conditions there was a clear cost of inbreeding to wing length; adults from inbred lines were much smaller than adults from large populations (females: F_1,82_ = 67.189, P < 0.001, males: F_1,82_ = 22.804, P < 0.001). There were substantial differences in wing length between replicate isofemale lines (females: F_4,45_ = 8.303, P < 0.001, males: F_4,45_ = 10.751, P < 0.001) and inbred lines (females: F_3,30_ = 20.703, P < 0.001, males: F_3,30_ = 9.729, P < 0.001) and smaller, but still significant, differences between females of the large (F_4,45_ = 3.102, P = 0.024) and small (F_4,44_ = 2.777, P = 0.038) populations. Wing lengths of adults from the large populations at F_13_ were smaller than those from the ancestral population at F_4/5_ (females: F_1,58_ = 10.472, P = 0.002, males: F_1,58_ = 10.519, P = 0.002) which could reflect adaptation to artificial rearing conditions. However, there were no differences between the laboratory and field populations from Cairns and Innisfail (all P > 0.05).

**Figure 6.**
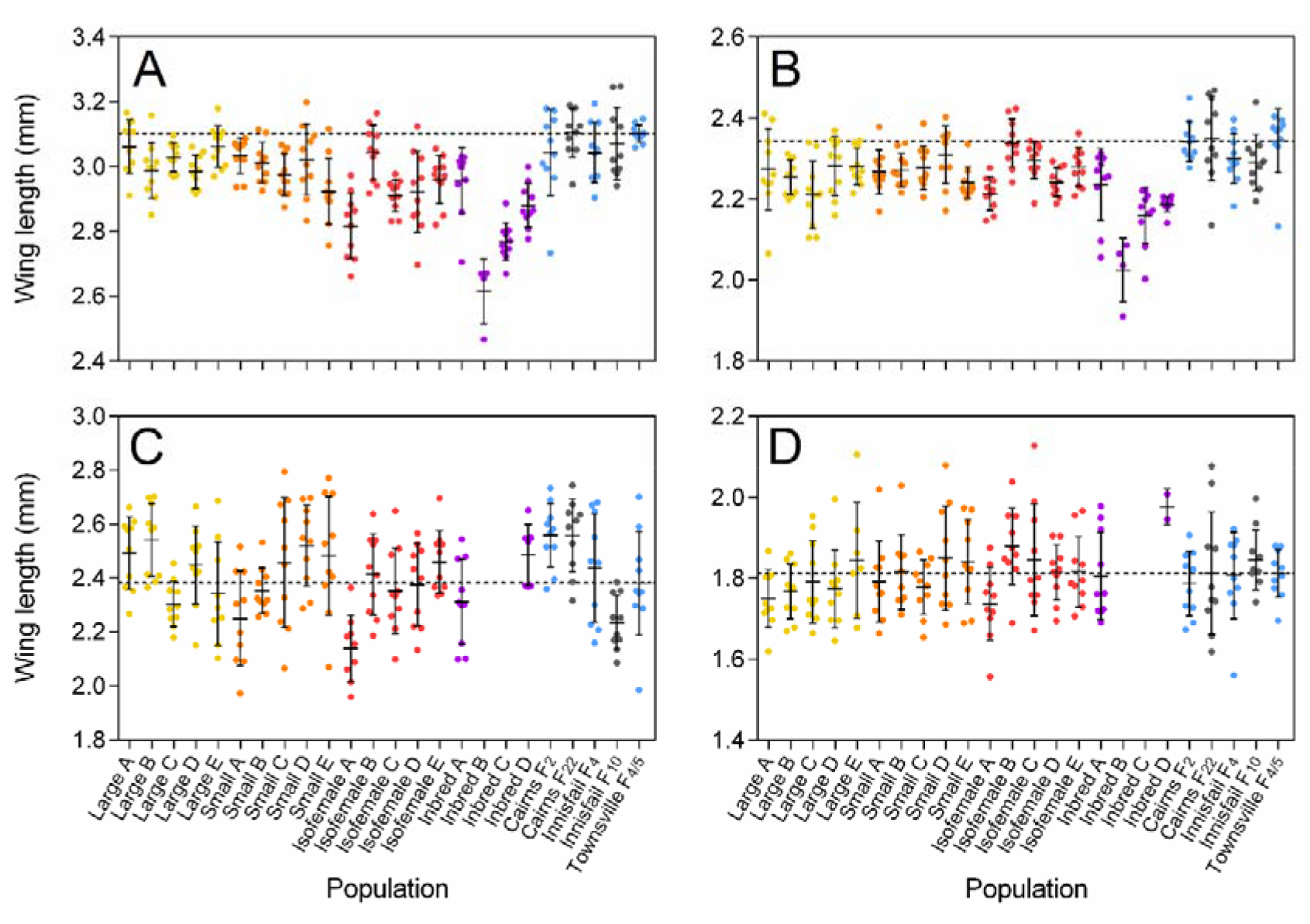
Wing length of *Aedes aegypti* F_13_ laboratory populations maintained at different census sizes. Wing lengths were measured from (A&C) female and (B&D) male adults when reared under (A&B) high nutrition (food provided *ad libitum)* and (C&D) low nutrition (0.1 mg of TetraMin^®^ per larva every 2 days) conditions. Replicates of the large populations (Townsville F_13_, census size 400) are in yellow, small populations (Townsville F_13_, census size 100) in orange, isofemale lines (Townsville F_13_) in red and inbred lines (Townsville F_13_) in purple. Other laboratory populations are shown in gray and ancestral / field populations in pale blue. Up to ten wings were measured for each group, though some inbred lines had less than 10 adults available, and several measurements were discarded due to damaged wings. Inbred lines B and C were not tested under low nutrition conditions. The dashed line represents the mean wing length of the Townsville F_4/5_ ancestral population. Error bars are standard deviations.

### Overall performance

We calculated an index of performance for each Townsville population at F_13_ relative to the ancestral population (Townsville F_4/5_) using the data for fecundity, egg hatch proportion, larval development time and survival to adulthood (under high nutrition conditions) available for each population (Figure 7). The large populations (census size 400) consistently performed better than the ancestral population (one sample t-test, P = 0.008) which indicates a positive effect of artificial rearing conditions on performance in the laboratory. The increased performance of laboratory populations was largely due to shorter larval development time (S3 Appendix). The relative performance of populations declined substantially with increasing levels of inbreeding; isofemale lines and inbred lines had much poorer performance than the ancestral population. This fitness deficit could largely be restored by crossing inbred mosquitoes to an outbred population (S4 Appendix). The Cairns laboratory population (F_22_) had increased performance over the field (F_2_) population (relative performance index: 1.080) but the Innisfail laboratory population (F_10_) had decreased performance over the field population (F_4_) (relative performance index: 0.960). Laboratory populations therefore did not always exhibit increased performance over the populations that were colonized more recently.

**Figure 7.**
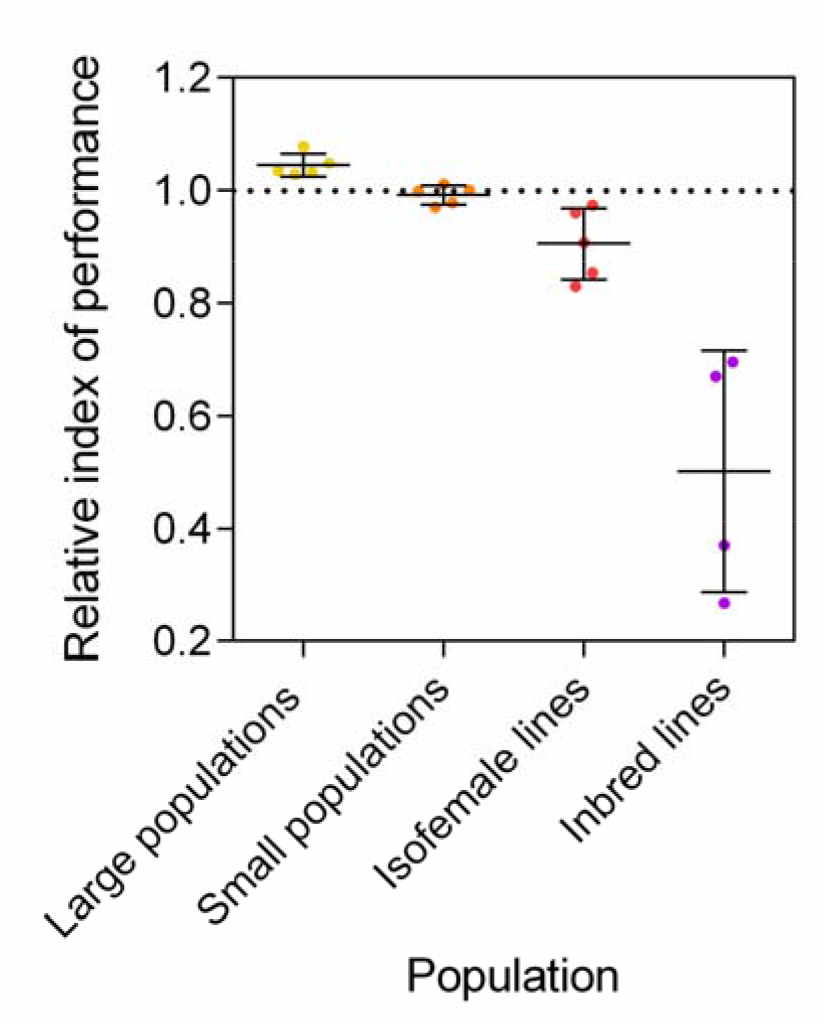
Relative performance of *Aedes aegypti* F_13_ laboratory populations maintained at different census sizes. Each data point represents the performance index of a single replicate population relative to the ancestral population (Townsville F_4/5_) which is represented by the black dotted line. Black bars indicate means and standard deviations.

### Effective population size

We estimated the effective population size (N_e_) of the replicate Townsville populations at F_13_ relative to the ancestral population (F_1_) using pooled RADseq and the Nest R package v1.1.9 [64]. The *N_e_*(*JR*) and *N_e_*(*P*) methods provided similar estimates of Ne but *N_e_*(*W*) provided estimates that were in many cases much larger than the census sizes. For estimates calculated using the *N_e_*(*JR*) and *N_e_*(*P*) methods, Ne declined substantially with decreasing census size (Table 1). Ratios of Ne to census size calculated using the *N_e_*(*P*) method were low, though the small populations (census size 100, mean N_e_/N = 0.250) had higher ratios than large populations (census size 400, mean N_e_/N = 0.143). The index of performance for each population increased dramatically with increasing N_e_ but levelled off at higher N_e_ (S1 Figure). These findings demonstrate a clear association between N_e_ and fitness (Spearman’s rank-order correlation: *ρ* = 0.973, *P* < 0.001, n = 17) but suggests that an N_e_ greater than used in the large populations will lead to only small fitness improvements.

**Table 1.**
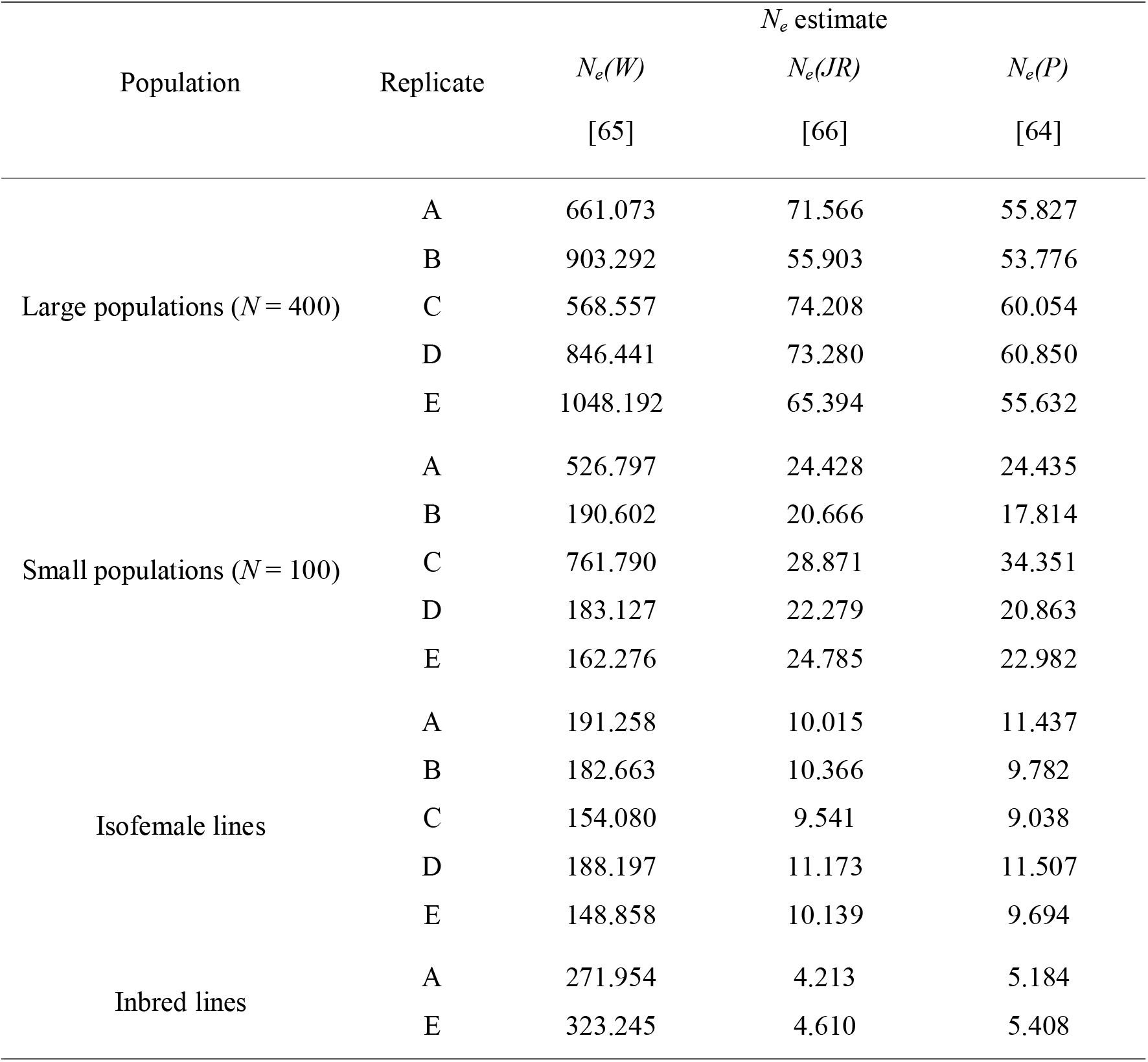
Effective population sizes (N_e_) of *Aedes aegypti* F_13_ laboratory populations maintained at different census sizes, calculated using three temporal methods.

| Population | Replicate | $N_e$ estimate | | |
| --- | --- | --- | --- | --- |
| | | $N_e(W)$ | $N_e(JR)$ | $N_e(P)$ |
|  |  | [65] | [66] | [64] |
| Large populations ($N = 400$) | A | 661.073 | 71.566 | 55.827 |
|  | B | 903.292 | 55.903 | 53.776 |
|  | C | 568.557 | 74.208 | 60.054 |
|  | D | 846.441 | 73.280 | 60.850 |
|  | E | 1048.192 | 65.394 | 55.632 |
| Small populations ($N = 100$) | A | 526.797 | 24.428 | 24.435 |
|  | B | 190.602 | 20.666 | 17.814 |
|  | C | 761.790 | 28.871 | 34.351 |
|  | D | 183.127 | 22.279 | 20.863 |
|  | E | 162.276 | 24.785 | 22.982 |
| Isofemale lines | A | 191.258 | 10.015 | 11.437 |
|  | B | 182.663 | 10.366 | 9.782 |
|  | C | 154.080 | 9.541 | 9.038 |
|  | D | 188.197 | 11.173 | 11.507 |
|  | E | 148.858 | 10.139 | 9.694 |
| Inbred lines | A | 271.954 | 4.213 | 5.184 |
|  | E | 323.245 | 4.610 | 5.408 |

### Mating competitiveness

The success of sterile and incompatible insect programs will depend on the ability of modified males to mate successfully in the wild. We tested to see if inbreeding and laboratory maintenance had any effect on male mating competitiveness. Males from the Cairns F_2_, F_7_ or F_27_ and inbred (Inbred A F_18_) populations competed for access to F2 females against a standard competitor infected with *Wolbachia* (*w*AlbB strain) (Figure 8). Hatch proportions did not differ significantly between the F_2_, F_7_ and F_27_ populations (one-way ANOVA: F_2,12_ = 0.829, P = 0.460), but were markedly reduced for inbred males relative to the other populations (one-way ANOVA: F_1,18_ = 39.784, P < 0.001). These results indicate that male mating success in laboratory cages is not affected by long-term laboratory maintenance, but can be decreased by inbreeding. The poor performance of inbred males was likely due to reduced mating success and not a paternal effect on female fertility, as crosses between inbred males and Cairns F_2_ females produced eggs with high hatch proportions (S4 Appendix).

**Figure 8.**
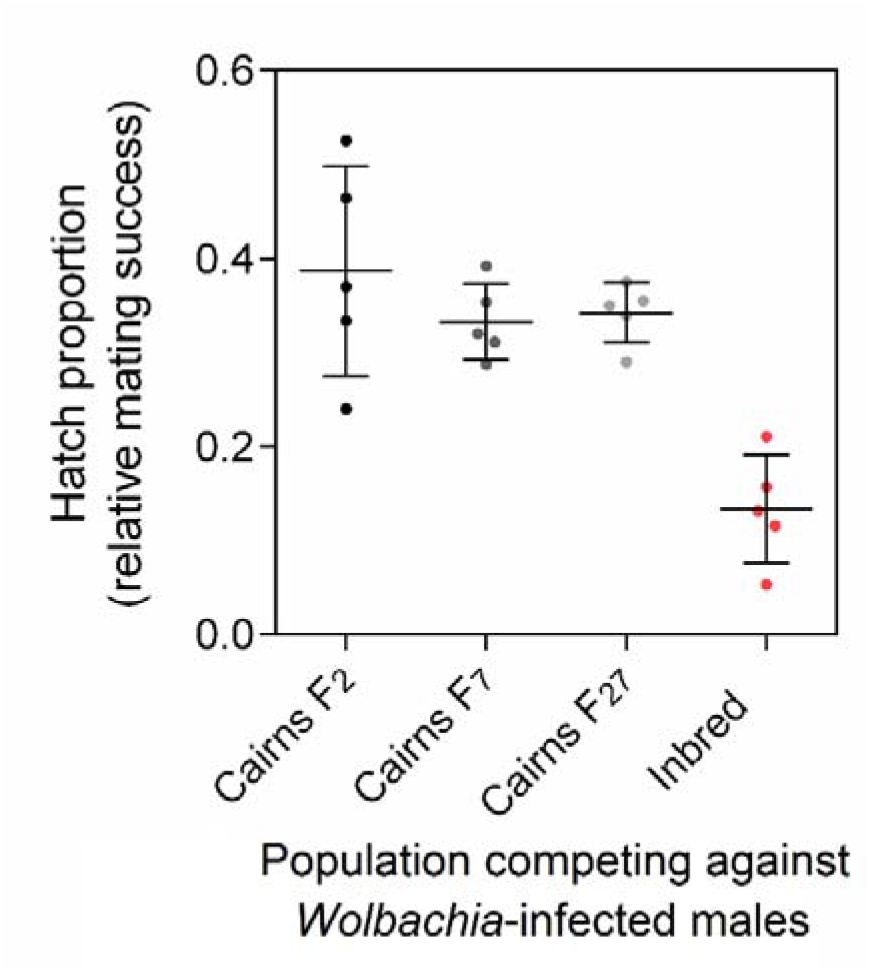
Relative mating success of *Aedes aegypti* males maintained in the laboratory for different numbers of generations. We tested the relative mating success of males from Cairns F_2_, F_7_ and F_27_ populations when competing against Wolbachia-infected males for access to Cairns F_2_ females. An inbred colony (Inbred A F_18_) was included for comparison. Higher hatch proportions indicate increased mating success of the experimental males relative to *Wolbachia*-infected males. Each data point represents the mean egg hatch proportion from a cage of 50 females. Black bars indicate means and standard deviations.

## Discussion

We performed a comprehensive assessment of inbreeding and laboratory adaptation in *Ae. aegypti* mosquitoes to inform rear and release programs for arbovirus control. Our study is the first to investigate the effects of inbreeding on *Ae. aegypti* fitness directly by comparing outbred and inbred lines derived from the same population, and the first that links fitness costs to reductions in effective population size as assessed through genomic markers. We look for evidence of adaptation by comparing laboratory populations to their direct ancestor concurrently and use replicate populations to separate fitness changes due to adaptation from drift and founder effects, two approaches which have not been previously applied in mosquitoes.

We find evidence of laboratory adaptation in colonized *Ae. aegypti* populations, but changes in trait means were small in magnitude and directions were often inconsistent between populations. All replicate laboratory populations from Townsville developed faster and were smaller than mosquitoes from the ancestral population. These changes could be a response to selection for abbreviated development in the laboratory, despite efforts to avoid selection in our laboratory rearing protocol [63]. Shorter developmental periods are often observed in laboratory-adapted insects [57], particularly under mass-rearing conditions that favor the rapid production of insects [67, 68]. In contrast, development times can increase in colonized *Drosophila* maintained with overlapping generations, where there is less selection against slow developing individuals [24].

When fitness traits were combined into an overall index of performance, we found that laboratory populations maintained at a large census size usually had greater performance than field populations. This finding is consistent with other insects, where fitness under laboratory conditions tends to improve with laboratory maintenance (Hoffmann and Ross unpublished). The rate of adaptation in our laboratory colonies of *Ae. aegypti* was slower than other mosquito species and insects in general (Hoffmann and Ross unpublished). *Ae. aegypti* collected from the field performed well from the first generation in the laboratory; rates of adaptation are likely to be higher for species such as *Ae. notoscriptus* where the laboratory environment is suboptimal and only a small proportion of individuals are able to reproduce in the initial generations [38]. Populations tested at F_2_ did not tend to differ from laboratory populations in terms of trait means, but some traits exhibited greater variation at F_2_. This suggests that *Ae. aegypti* could lose variation with laboratory maintenance, though other traits for other populations at F2 had similar variances to laboratory populations. Our main comparisons were between populations at F_4/5_ and F_13_; if substantial laboratory adaptation occurs then we would be unable to detect it with these comparisons. Our population comparisons could also be confounded by selection on the ancestral population due to eggs experiencing quiescence (Townsville and Innisfail populations) or differences present in populations collected from the same location but at different times (Cairns populations). Other factors such as gut microbiota could also confound our comparisons between laboratory and field populations because the microbiome can greatly influence mosquito life history traits [69, 70]. Gut microbiota are much less diverse in colonized mosquitoes [71] and tend to be similar in laboratory populations regardless of geographic origin [72]. This could be an issue when comparing field and laboratory populations.

Few studies on laboratory adaptation in insects attempt to separate the effects of laboratory adaptation from drift or founder effects (Hoffmann and others [30] is one exception) which are likely to be substantial when establishing small laboratory colonies. We used replicate populations to avoid this issue; consistent divergence in colonized populations from the ancestral population indicates adaptation, variation between replicate populations immediately after establishment indicates founder effects, and divergence between replicate populations at the time indicates drift. We found that replicate populations at the same census size differ significantly from each other for several fitness traits, both at F_5_ and at F_13_, particularly for populations maintained at low census sizes. Fitness differences between replicate populations were not always consistent between F_5_ and F_13_, suggesting that both founder effects and drift occur. These findings are of concern for laboratory studies that compare traits between populations maintained separately. Researchers should consider using replicate populations when conducting experiments or outcross populations frequently to maintain similar genetic backgrounds [73].

We demonstrate that inbreeding is extremely costly to *Ae. aegypti* fitness. Most inbred lines were lost across the experiment, and the remaining lines performed substantially worse than outbred populations. Relatively few studies have specifically addressed the effects of inbreeding on mosquito fitness. Powell and Evans [74] observed that inbreeding *Ae. aegypti* through full-sib mating reduces heterozygosity by much less than expected based on theory, and deleterious recessive alleles must therefore be common. Koenraadt and others [75] reported fitness costs of inbred *Ae. aegypti* larvae relative to a wild population, and inbreeding through full-sib mating reduces fitness in other *Aedes* species [76, 77]. We demonstrate that mosquito populations inbred intentionally, for instance, to generate homozygous transgenic strains [78, 79], will likely suffer from severe inbreeding depression. However, it may be possible to retain partial fitness if there is also selection for certain life history traits during inbreeding [80]. We show that laboratory populations maintained at low census sizes (N = 100) also experience inbreeding depression, and the loss of fitness correlates strongly with decreased effective population size. Thus, laboratories should ensure that population sizes in colonized mosquitoes are sufficiently high to maintain their fitness. Our laboratory populations for these experiments were each established from only a few hundred individuals, and we would recommend that larger numbers be used to avoid bottlenecks.

Our laboratory populations at F_13_ had a substantially lower N_e_ than field populations from Townsville [81] and other locations around the world [82]. However, ratios of Ne to census size (N_e_/N) were similar to ratios reported in nature [82]. N_e_/N ratios were larger in the small laboratory populations (N = 100) than in the large ones (N = 400), consistent with a study of *Drosophila* populations maintained at different census sizes [83]. Low N_e_/N ratios indicate that reproductive success varies greatly between individuals [84, 85] and this appears to be the case for large colonized populations of *Ae. aegypti*. Unequal contributions to the next generation occur because we sample only a few hundred individuals randomly from a pool of thousands of larvae, and we do not equalize offspring from each female to establish the next generation [63].

We demonstrate that the consequences of laboratory maintenance in *Ae. aegypti* can be minimized by maintaining large population sizes, but there are several other ways to maintain the fitness of colonized mosquito populations. The simplest approach is to cross laboratory colonies to an outbred population [73]. Gene flow into inbred populations commonly leads to a fitness improvement [86] and we also show that the fitness of inbred *Ae. aegypti* can be improved through a single generation of outcrossing. Increased performance of hybrids has been demonstrated in *Anopheles* mosquitoes [87–89] and the Queensland fruit fly [90], with fitness improvements in the F_1_. Crosses between different laboratory lines can also be used to determine whether changes in fitness in laboratory populations are due to inbreeding or adaptation [89]. Rates of laboratory adaptation can be slowed by using more natural rearing environments. Knop and others [50] compared two methods of rearing *Culex tarsalis* and found that colonies maintained in larger cages at a variable temperature and more complex environmental conditions had a slower rate of laboratory adaptation. Ng’habi and others [56] found that rearing *Anopheles arabiensis* under semi-field conditions preserved their similarity to the wild population and reduced the extent of inbreeding. Quality control methods such as screening mosquitoes for their flight capacity can also be used to increase fitness before their deployment for disease control programs [91].

In summary, we provide evidence for inbreeding depression effects and a small effective population size relative to census size in laboratory mosquito populations, along with some limited laboratory adaptation particularly in large populations. Our results have implications for the maintenance of insects in the laboratory, particularly for those destined for open field releases. While we find that life history traits of *Ae. aegypti* do not change consistently with laboratory maintenance, traits where selective pressures are absent in the laboratory, such as flight ability, feeding behaviour and thermal tolerance might still be compromised.

## Materials and methods

### Replicate population establishment

*Aedes* eggs collected from ovitraps near Townsville, Australia, in September 2015 [92] were hatched and reared in the laboratory (see *Colony maintenance*). *Ae. aegypti* larvae were separated from other species based on an identification key [93]. A total of 327 *Ae. aegypti* adults (171 males and 156 females) were obtained and added to a single 19.7 L colony cage. Females were blood-fed, and all eggs laid were pooled and hatched in a single tray containing 3 L of water. Larvae were selected at random and divided into groups to establish replicate populations (Figure 1). Five populations were maintained at a census size of 400 adults (large populations) and five populations were maintained at a census size of 100 adults (small populations). Twenty adult females were also isolated for oviposition. The offspring from five isolated females were used to establish five additional populations maintained at a census size of 100 adults (isofemale lines), while the offspring from ten females were maintained with a single male and female each (inbred lines). At least two mating pairs were established for each inbred line (S2 Table), but only a single pair was used to found the next generation. Their offspring underwent full-sib mating each generation for nine generations, then all progeny were interbred during F_12_ to build up numbers for experiments. All replicate populations were maintained until F_13_ when experimental comparisons were performed. All adults from the ancestral Townsville population (F_1_) and the replicate populations at F_13_ were stored in absolute ethanol at −20°C for pooled double-digest RADseq. Only two inbred lines had sufficient numbers for RADseq due to the loss of most inbred lines over the course of full-sib mating (S2 Table).

*Ae. aegypti* eggs can withstand desiccation and remain viable for up to one year [43]. We utilised this ability to perform direct comparisons between the ancestral population and the derived populations simultaneously. Eggs laid by F_2_ and F_3_ females were stored under humid conditions for several months at 26°C and then hatched at the same time as eggs laid by F_11_ females from the other populations. A colony derived from larvae that hatched was maintained under standard conditions for one generation and their progeny (F_4/5_) were used for experiments alongside the populations at F_13_. Colonies derived from eggs collected from Cairns and Innisfail, Australia were also used for experimental comparisons. These colonies were maintained as single caged populations with a census size of 400 individuals. Quiescent eggs from the Innisfail population were also used to generate a colony that had experienced fewer generations of maintenance under laboratory conditions. Eggs collected from Cairns at a later stage were used to establish a colony for comparisons with the Cairns colony at F_22_.

### Colony maintenance

All populations were maintained in a controlled temperature laboratory environment (26 ± 0.5°C and 50-70% relative humidity, with a 12:12 h light:dark photoperiod) following the protocol described by Ross and others [63]. This protocol is designed to reduce selection against individuals that are slow or quick to develop, mature, mate, blood feed, oviposit or hatch, and to minimize mortality at each life stage. To maintain each population, all eggs from the previous generation were pooled and a random subset of larvae was provided with food (TetraMin® tropical fish food tablets, Tetra, Melle, Germany) *ad libitum* and reared to adulthood. For the large populations, 400 adults were selected at random and added to 19.7 L cages, while for the small populations and isofemale lines, 100 adults were added to 12 L cages. For the inbred lines, a single male and female were added to a 1.5 L cage. Except for the inbred lines, sex ratios were maintained naturally, and equal numbers of males and females were not counted. All cages were provided with a source of water and 10% sucrose. Approximately three days after the last adult had emerged, females were blood fed on a single human volunteer. Two days after blood feeding, cups containing larval rearing water and lined with sandpaper strips were introduced into the cages. Eggs laid on the sandpaper strips were collected over the span of one week, and all eggs were hatched three days after females had ceased oviposition. We followed this procedure until the Townsville populations were at F_13_, with each generation taking 28 days to complete. Blood feeding of mosquitoes on human subjects was approved by the University of Melbourne Human Ethics Committee (approval #: 0723847). All volunteers provided informed written consent.

### Fitness comparisons between Townsville F_13_ populations

We compared all Townsville populations at F_13_ for their development time, survival to adulthood and wing length under two nutrition conditions, and the fecundity and egg hatch rate of females reared under high nutrition conditions. Not all inbred lines were included in the experiments as the majority were lost by F_13_ (S2 Table). Two of the four remaining inbred lines were only tested under high nutrition conditions due to low numbers, and these lines later became extinct (S2 Table). Cairns (F_2_ and F_22_), Innisfail (F_4_ and F_10_) and Townsville (F_4/5_) populations were included in all experiments.

One hundred larvae from each population were reared in containers with 500 mL of water and provided with TetraMin^®^ *ad libitum* (high nutrition) or with 0.1 mg of TetraMin^®^ per larva every 2 days (low nutrition). Four replicate containers were reared for each population, except for two inbred lines where less than 400 larvae were obtained. A random subset of females from each population that emerged from the high nutrition treatment were blood fed and then isolated for oviposition. Eggs collected from each female were counted and hatched three days post-collection. Egg hatch rates were determined by calculating the proportion of eggs that had a detached cap. Isolated females were blood fed again after one week and fecundity and egg hatch rate were measured for a second gonotrophic cycle. Wings from 10 males and 10 females selected at random from each population and each nutrition treatment were dissected and measured for their length according to methods described previously [94].

Fitness data from the Townsville populations at F_13_ were used to estimate the performance of each population relative to the Townsville F_4/5_ ancestral population. We simplified an equation from Livdahl and Sugihara [95] to calculate performance from fecundity, egg hatch, survival and larval development time data. *F* is the mean fecundity of each population multiplied by egg hatch proportion, *S* is the mean proportion of larvae surviving to adulthood, and *D* is the mean larval development time in days. The performance index of each population at F_13_ was divided by the performance index of the ancestral population to determine their relative performance.

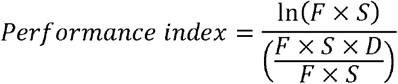

### Male mating competitiveness

We tested the male mating competitiveness of populations from Cairns that were at F_2_, F_7_ or F_27_ in the laboratory, and an inbred line from Townsville (Inbred A) at F_18_. Males from all populations competed against males infected with the *w*AlbB strain of *Wolbachia* for access to F_2_ females in a caged laboratory environment. Males infected with *w*AlbB induce complete sterility (eggs do not hatch) when crossed to uninfected females under standard laboratory conditions [12, 96]. Thus, the competitive ability of each population relative to *w*AlbB-infected males can be estimated by scoring egg hatch rate from crosses between uninfected females and a mix of Wolbachia-infected and uninfected males [36, 97]. We established 12 L cages containing 25 males from each population (F_2_, F_7_, F_27_ or inbred) and 25 males infected with *w*AlbB. Five replicate cages were established for each treatment. We then aspirated ten Cairns F_2_ females into each cage. This was repeated five times at 1 hr intervals, for a total for 50 females per cage. Staggered releases were chosen to increase the level of male-male competition; adding all females to a cage at once would not provide many males with an opportunity to inseminate multiple females. All individuals were reared under the same conditions for this experiment (see *Colony maintenance*), and males were at least 24 hr old, and females at least 48 hr old before the sexes were combined. Females were blood fed three days after mating and a single cup filled with larval rearing water and lined with a sandpaper strip was added to each cage. Sandpaper strips were collected daily and photographed, and the number of eggs on each strip was counted in ImageJ [98] using the Cell Counter plugin (https://imagej.nih.gov/ij/plugins/cell-counter.html). Eggs were hatched three days post-collection and larvae were counted three days after hatching. Egg hatch rates were estimated by dividing the number of larvae counted by the number of eggs from each cage.

### Pooled double-digest RADseq library preparation

We used pooled double-digest RADseq to determine the effective population size (N_e_) of the seventeen replicate populations from Townsville at F_13_ relative to their ancestral population (F_1_). These included the five large populations, five small populations, five isofemale lines and two inbred lines. We prepared a library following methods described by Rašić and others [99] and Schmidt and others [100], but modified the protocol for pooled mosquitoes. DNA was extracted from four pools of 20 adult mosquito heads from each population, with two pools for each sex, using a Roche DNA Isolation Kit for Cells and Tissues (Roche, Pleasanton, CA, USA). DNA from each pool was quantified using a Qubit dsDNA HS assay kit and a Qubit 2.0 fluorometer (Thermo Fisher Scientific, Life Technologies Holdings Pte Ltd Singapore) and the four pools for each population were combined after a normalization step.

750 ng of DNA from each population was digested in a 50 μL reaction with 20 units each of *Eco-R1-HF* and *SphI-HF* restriction enzymes (New England Biolabs, Beverly, MA, USA), NEB CutSmart^®^ buffer and water for three hours at 37°C with no heat kill step. Restriction enzymes that cut less frequently were chosen to produce fewer SNPs while providing more coverage. The digestion products were cleaned with 75 μL of Ampure XP™ paramagnetic beads (Beckman Coulter, Brea, CA) and then ligated with modified Illumina P1 and P2 adapters overnight at 16°C with 1000 units of T4 ligase and 1× T4 buffer (New England Biolabs, Beverly, MA, USA) in a 45 μL reaction volume, then heat deactivated for 10 minutes at 65°C. Ligations were cleaned using 75 μL of paramagnetic beads and adapter-ligated DNA fragments from all eighteen populations were pooled. We then used a Pippin-Prep 2% gel cassette (Sage Sciences, Beverly, MA, USA) to select fragments with a size range of 350-450 bp. The final library was generated by pooling 38 10 μL PCR reactions and run for 12 cycles; each reaction contained 5 μL of Phusion High Fidelity 2× Master mix (New England Biolabs, Beverly MA, USA), 2 μL each of 10 μM forward and reverse Illumina primers and 2 μL of size-selected DNA. The pooled PCR reactions were cleaned with 300 μL of paramagnetic beads, and a single library with DNA from 1440 *Ae. aegypti* from the eighteen populations was sequenced in a single Illumina HiSeq 2000 lane to obtain 100 bp paired-end reads.

### Data processing and effective population size estimates

We checked the quality of the raw sequencing data with FastQC v0.11.5 [101] and then used the process_radtags component of Stacks v1.46 [102] to demultiplex the populations, allowing for one mismatch. Reads were trimmed to 80 bp and then aligned to the *Aedes aegypti* reference genome AaegL4 [103] using bowtie2 v2.3.0 [104]. Ambiguous alignments (minimum mapping quality below 20) were discarded and alignments were converted to SAM format and sorted using SAMtools v1.4 [105]. Sorted files were then converted to mpileup format, with each file containing the ancestral population and one of the seventeen derived populations. These files were converted to sync format using the mpileup2sync.jar tool from Popoolation2 [106]. We then estimated effective population size (N_e_) using the Nest R package v1.1.9 with three different methods [64].

### Statistics on life history traits

All data were analyzed using SPSS statistics version 24.0 for Windows (SPSS Inc, Chicago, IL). All data were checked for normality using Shapiro-Wilk tests. Data that were normally distributed were analyzed using ANOVA and data that were not normal were analyzed with Kruskal-Wallis and Mann-Whitney U tests.

## Acknowledgements

The authors thank Scott Ritchie’s group at James Cook University for providing field-collected eggs used in the experiments. We thank Elizabeth Valerie, Shani Wong, Fionna Zhu and Isabelle Foo for their assistance with mosquito colony maintenance and fitness experiments. We also thank Peter Kriesner, Tom Schmidt, Moshe Jasper, Gordana Rašić and Pip Griffin for their advice on the design and analysis of pooled RADseq. PAR was supported by an Australian Government Research Training Program Scholarship. Funding was provided by the National Health and Medical Research Council via their Program and Fellowship schemes.

## Supporting information

**S1 Table. Studies of laboratory adaptation in mosquitoes**. Studies compared mosquito populations that were maintained in the laboratory for different numbers of generations. Focal traits were either tested over successive generations in the laboratory, or field and laboratory populations were compared simultaneously.

**S2 Table. Maintenance and loss of inbred *Aedes aegypti* populations during successive generations of full-sib mating**. The number of replicate male and female pairs for each line increased as inbred lines were lost.

**S1 Figure. The effective population size (N_e_) of *Aedes aegypti* populations at F_13_ *versus* their index of performance**. Large populations (Townsville F_13_, census size 400) are in yellow, small populations (Townsville F_13_, census size 100) in orange, isofemale lines (Townsville F_13_) in red and inbred lines (Townsville F_13_) in purple. The relative performance of the Townsville F_4/5_ population is represented by the black dotted line. Effective population size was calculated using the *N_e_*(*P*) method.

**S1 Appendix. Pilot experiments comparing fitness differences between Townsville F_2_ and Cairns F_11_ *Aedes aegypti* populations**.

**S2 Appendix. Fitness comparisons between large *Aedes aegypti* populations (census size 400) and inbred lines from Townsville at F_5_**.

**S3 Appendix. The extent of laboratory adaptation in *Aedes aegypti* and comparisons with other insects**.

**S4 Appendix. Outcrossing of an inbred *Aedes aegypti* population**.

**S5 Appendix. Larval competitiveness of *Aedes aegypti* laboratory populations**.

## References

1. Benedict M, Robinson AS. The first releases of transgenic mosquitoes: an argument for the sterile insect technique. Trends in Parasitology. 2003;19(8):349–55. doi: 10.1016/s1471-4922(03)00144-2.

2. Bellini R, Calvitti M, Medici A, Carrieri M, Celli G, Maini S. Use of the sterile insect technique against *Aedes albopictus* in Italy: first results of a pilot trial. Area-Wide Control of Insect Pests. 2007:505–15.

3. Bellini R, Medici A, Puggioli A, Balestrino F, Carrieri M. Pilot field trials with *Aedes albopictus* irradiated sterile males in Italian urban areas. Journal of Medical Entomology. 2013;50(2):317–25. doi: 10.1603/me12048.

4. McGraw EA, O'Neill SL. Beyond insecticides: new thinking on an ancient problem. Nature reviews Microbiology. 2013;11(3):181–93. doi: 10.1038/nrmicro2968. PubMed PMID: 23411863.

5. Ritchie SA, Johnson BJ. Advances in vector control science: rear-and-release strategies show promise.but don't forget the basics. J Infect Dis. 2017;215(suppl_2):S103–S8. doi: 10.1093/infdis/jiw575. PubMed PMID: 28403439.

6. Harris AF, McKemey AR, Nimmo D, Curtis Z, Black I, Morgan SA, et al. Successful suppression of a field mosquito population by sustained release of engineered male mosquitoes. Nature biotechnology. 2012;30(9):828–30. doi: 10.1038/nbt.2350. PubMed PMID: 22965050.

7. Lacroix R, McKemey AR, Raduan N, Kwee Wee L, Hong Ming W, Guat Ney T, et al. Open field release of genetically engineered sterile male *Aedes aegypti* in Malaysia. PloS one. 2012;7(8):e42771. doi: 10.1371/journal.pone.0042771. PubMed PMID: 22970102; PubMed Central PMCID: PMCPMC3428326.

8. Carvalho DO, McKemey AR, Garziera L, Lacroix R, Donnelly CA, Alphey L, et al. Suppression of a field population of *Aedes aegypti* in Brazil by sustained release of transgenic male mosquitoes. PLoS neglected tropical diseases. 2015;9(7):e0003864. doi: 10.1371/journal.pntd.0003864. PubMed PMID: 26135160.

9. Garziera L, Pedrosa MC, de Souza FA, Gómez M, Moreira MB, Virginio JF, et al. Effect of interruption of over-flooding releases of transgenic mosquitoes over wild population of *Aedes aegypti*: two case studies in Brazil. Entomologia Experimentalis et Applicata. 2017;164(3):327–39. doi: 10.1111/eea.12618.

10. Curtis Z, Matzen K, Neira Oviedo M, Nimmo D, Gray P, Winskill P, et al. Assessment of the impact of potential tetracycline exposure on the phenotype of *Aedes aegypti* OX513A: implications for field use. PLoS neglected tropical diseases. 2015;9(8):e0003999. doi: 10.1371/journal.pntd.0003999. PubMed PMID: 26270533; PubMed Central PMCID: PMC4535858.

11. Ferguson NM, Hue Kien DT, Clapham H, Aguas R, Trung VT, Bich Chau TN, et al. Modeling the impact on virus transmission of *Wolbachia*-mediated blocking of dengue virus infection of *Aedes aegypti*. Sci Transl Med. 2015;7(279):279ra37.

12. Xi Z, Khoo CC, Dobson SL. *Wolbachia* establishment and invasion in an *Aedes aegypti* laboratory population. Science. 2005;310(5746):326–8. doi: 10.1126/science.1117607. PubMed PMID: 16224027.

13. Walker T, Johnson PH, Moreira LA, Iturbe-Ormaetxe I, Frentiu FD, McMeniman CJ, et al. The wMel *Wolbachia* strain blocks dengue and invades caged *Aedes aegypti* populations. Nature. 2011;476(7361):450–3. PubMed PMID: 21866159.

14. O'Connor L, Plichart C, Sang AC, Brelsfoard CL, Bossin HC, Dobson SL. Open release of male mosquitoes infected with a *Wolbachia* biopesticide: field performance and infection containment. PLoS neglected tropical diseases. 2012;6(11):e1797. doi: 10.1371/journal.pntd.0001797. PubMed PMID: 23166845; PubMed Central PMCID: PMC3499408.

15. Mains JW, Brelsfoard CL, Rose RI, Dobson SL. Female adult *Aedes albopictus* suppression by *Wolbachia*-infected male mosquitoes. Scientific reports. 2016;6:33846. doi: 10.1038/srep33846. PubMed PMID: 27659038; PubMed Central PMCID: PMCPMC5034338.

16. Hoffmann AA, Montgomery BL, Popovici J, Iturbe-Ormaetxe I, Johnson PH, Muzzi F, et al. Successful establishment of *Wolbachia* in Aedes populations to suppress dengue transmission. Nature. 2011;476(7361):454–7. doi: 10.1038/nature10356. PubMed PMID: 21866160.

17. Nguyen TH, Nguyen HL, Nguyen TY, Vu SN, Tran ND, Le TN, et al. Field evaluation of the establishment potential of wMelPop *Wolbachia* in Australia and Vietnam for dengue control. Parasites & vectors. 2015;8:563. doi: 10.1186/s13071-015-1174-x. PubMed PMID: 26510523; PubMed Central PMCID: PMC4625535.

18. Schmidt TL, Barton NH, Rasic G, Turley AP, Montgomery BL, Iturbe-Ormaetxe I, et al. Local introduction and heterogeneous spatial spread of dengue-suppressing *Wolbachia* through an urban population of *Aedes aegypti*. PLoS biology. 2017;15(5):e2001894. doi: 10.1371/journal.pbio.2001894. PubMed PMID: 28557993.

19. Winskill P, Harris AF, Morgan SA, Stevenson J, Raduan N, Alphey L, et al. Genetic control of *Aedes aegypti*: data-driven modelling to assess the effect of releasing different life stages and the potential for long-term suppression. Parasites & vectors. 2014;7(1):68.

20. Leftwich PT, Bolton M, Chapman T. Evolutionary biology and genetic techniques for insect control. Evol Appl. 2015: 10.1111/eva.12280. doi: 10.1111/eva.12280.

21. Benedict MQ. Care and maintenance of anopheline mosquito colonies. The molecular biology of insect disease vectors: Springer; 1997. p. 3–12.

22. Munstermann LE. Care and maintenance of *Aedes* mosquito colonies. The Molecular Biology of Insect Disease Vectors: Springer; 1997. p. 13–20.

23. Carvalho DO, Nimmo D, Naish N, McKemey AR, Gray P, Wilke AB, et al. Mass production of genetically modified *Aedes aegypti* for field releases in Brazil. Journal of visualized experiments: JoVE. 2014;(83):e3579. doi: 10.3791/3579. PubMed PMID: 24430003; PubMed Central PMCID: PMC4063546.

24. Sgro CM, Partridge L. Evolutionary responses of the life history of wild-caught Drosophila melanogaster to two standard methods of laboratory culture. Am Nat. 2000;156(4):341–53. doi: Doi 10.1086/303394. PubMed PMID: WOS:000089408200002.

25. Simoes P, Santos J, Matos M. Experimental Evolutionary Domestication. Experimental evolution: Concepts, methods, and applications of selection experiments. 2009:89–110. doi: BOOK_DOI 10.1525/california/9780520247666.001.0001. PubMed PMID: WOS:000295511600005.

26. Reisen WK, Knop NF, Peloquin JJ. Swarming and mating behavior of laboratory and field strains of Culex tarsalis (Diptera: Culicidae). Annals of the Entomological Society of America. 1985;78(5):667–73.

27. Pereira R, Silva N, Quintal C, Abreu R, Andrade J, Dantas L. Sexual performance of mass reared and wild Mediterranean fruit flies (Diptera: Tephritidae) from various origins of the Madeira Islands. Florida Entomologist. 2007;90(1):10–4. doi: 10.1653/0015-4040(2007)90[10:spomra]2.0.co;2.

28. Rull J, Brunel O, Mendez ME. Mass rearing history negatively affects mating success of male Anastrepha ludens (Diptera: Tephritidae) reared for sterile insect technique programs. Journal of Economic Entomology. 2005;98(5):1510–6. Epub 2005/12/13. PubMed PMID: 16334318.

29. Bryant EH, Reed DH. Fitness decline under relaxed selection in captive populations. Conservation Biology. 1999;13(3):665–9.

30. Hoffmann AA, Hallas R, Sinclair C, Partridge L. Rapid loss of stress resistance in Drosophila melanogaster under adaptation to laboratory culture. Evolution. 2001;55(2):436–8. Epub 2001/04/20. PubMed PMID: 11308098.

31. Pimentel D, Schwardt H, Dewey J. Development and loss of insecticide resistance in the house fly. Journal of Economic Entomology. 1953;46(2):295–8.

32. Briscoe D, Malpica J, Robertson A, Smith GJ, Frankham R, Banks R, et al. Rapid loss of genetic variation in large captive populations of Drosophila flies: implications for the genetic management of captive populations. Conservation Biology. 1992;6(3):416–25.

33. Frankham R. Genetic adaptation to captivity in species conservation programs. Molecular ecology. 2008;17(1):325–33. doi: 10.1111/j.1365-294X.2007.03399.x. PubMed PMID: 18173504.

34. Reisen W, Milby M, Asman S, Bock M, Meyer R, McDonald P, et al. Attempted suppression of a semi-isolated Culex tarsalis population by the release of irradiated males: a second experiment using males from a recently colonized strain. Mosquito news. 1982;42(4):565–75.

35. Helinski ME, Harrington LC. Considerations for male fitness in successful genetic vector control programs. Ecology of parasite-vector interactions: Springer; 2013. p. 221–44.

36. Chambers EW, Hapairai L, Peel BA, Bossin H, Dobson SL. Male mating competitiveness of a *Wolbachia*-introgressed *Aedes polynesiensis* strain under semi-field conditions. PLoS neglected tropical diseases. 2011;5(8):e1271. doi: 10.1371/journal.pntd.0001271. PubMed PMID: 21829750; PubMed Central PMCID: PMC3149012.

37. Harris AF, Nimmo D, McKemey AR, Kelly N, Scaife S, Donnelly CA, et al. Field performance of engineered male mosquitoes. Nature biotechnology. 2011;29(11):1034–7. doi: 10.1038/nbt.2019. PubMed PMID: 22037376.

38. Watson TM, Marshall K, Kay BH. Colonization and laboratory biology of *Aedes notoscriptus* from Brisbane, Australia. J Am Mosq Control Assoc. 2000;16(2):138–42.

39. Oliva CF, Benedict MQ, Lempérière G, Gilles J. Laboratory selection for an accelerated mosquito sexual development rate. Malaria journal. 2011;10(1):135.

40. Huho BJ, Ng'habi KR, Killeen GF, Nkwengulila G, Knols BG, Ferguson HM. Nature beats nurture: a case study of the physiological fitness of free-living and laboratory-reared male Anopheles gambiae s.l. The Journal of experimental biology. 2007;210(Pt 16):2939–47. Epub 2007/08/11. doi: 10.1242/jeb.005033. PubMed PMID: 17690243.

41. Ponlawat A, Harrington LC. Age and body size influence male sperm capacity of the dengue vector *Aedes aegypti* (Diptera: Culicidae). J Med Ent. 2007;44(3):422–6.

42. Hassan MaM, El-Motasim WM, Ahmed RT, El-Sayed BB. Prolonged colonisation, irradiation, and transportation do not impede mating vigour and competitiveness of male Anopheles arabiensis mosquitoes under semi-field conditions in Northern Sudan. 2010.

43. Faull KJ, Williams CR. Intraspecific variation in desiccation survival time of *Aedes aegypti* (L.) mosquito eggs of Australian origin. J Vector Ecol. 2015;40(2):292–300.

44. Jong ZW, Kassim NFA, Naziri MA, Webb CE. The effect of inbreeding and larval feeding regime on immature development of *Aedes albopictus*. Journal of Vector Ecology. 2017;42(1):105–12. Epub 2017/05/16. doi: 10.1111/jvec.12244. PubMed PMID: 28504428.

45. Chadee DD, Beier JC. Factors influencing the duration of blood-feeding by laboratory-reared and wild *Aedes aegypti* (Diptera: Culicidae) from Trinidad, West Indies. Annals of Tropical Medicine and Parasitology. 1997;91(2):199–207. Epub 1997/03/01. PubMed PMID: 9307662.

46. Chadee DD, Beier JC, Mohammed RT. Fast and slow blood-feeding durations of *Aedes aegypti* mosquitoes in Trinidad. Journal of Vector Ecology. 2002;27(2):172–7. Epub 2003/01/28. PubMed PMID: 12546453.

47. Yeap HL, Endersby NM, Johnson PH, Ritchie SA, Hoffmann AA. Body size and wing shape measurements as quality indicators of *Aedes aegypti* mosquitoes destined for field release. The American journal of tropical medicine and hygiene. 2013;89(1):78–92. PubMed PMID: 23716403; PubMed Central PMCID: PMC3748492.

48. Allgood DW, Yee DA. Oviposition preference and offspring performance in container breeding mosquitoes: evaluating the effects of organic compounds and laboratory colonisation. Ecological Entomology. 2017. doi: 10.1111/een.12412.

49. Lima JB, Valle D, Peixoto AA. Adaptation of a South American malaria vector to laboratory colonization suggests faster-male evolution for mating ability. BMC Evolutionary Biology. 2004;4(1):12. Epub 2004/05/11. doi: 10.1186/1471-2148-4-12. PubMed PMID: 15132759; PubMed Central PMCID: PMCPMC420237.

50. Knop NF, Asman SM, Reisen WK, Milby MM. Changes in the biology of Culex tarsalis (Diptera: Culicidae) associated with colonization under contrasting regimes. Environmental entomology. 1987;16(2):405–14.

51. Haeger JS, O'Meara GF. Rapid incorporation of wild genotypes of Culex nigripalpus (Diptera: Culicidae) into laboratory-adapted strains. Annals of the Entomological Society of America. 1970;63(5):1390–1.

52. Lorenz L, Beaty BJ, Aitken THG, Wallis GP, Tabachnick WJ. The effect of colonization upon *Aedes aegypti* - susceptibility to oral infection with yellow fever virus. American Journal of Tropical Medicine and Hygiene. 1984;33(4):690–4. PubMed PMID: WOS:A1984TD91100028.

53. Salazar MI, Richardson JH, Sanchez-Vargas I, Olson KE, Beaty BJ. Dengue virus type 2: replication and tropisms in orally infected *Aedes aegypti* mosquitoes. BMC microbiology. 2007;7:9. doi: 10.1186/1471-2180-7-9. PubMed PMID: 17263893; PubMed Central PMCID: PMC1797809.

54. Vazeille M, Rosen L, Mousson L, Failloux A-B. Low oral receptivity for dengue type 2 viruses of *Aedes albopictus* from Southeast Asia compared with that of *Aedes aegypti*. The American journal of tropical medicine and hygiene. 2003;68(2):203–8.

55. Grimstad PR, Craig Jr GB, Ross QE, Yuill TM. *Aedes triseriatus* and La Crosse virus: geographic variation in vector susceptibility and ability to transmit. The American journal of tropical medicine and hygiene. 1977;26(5):990–6.

56. Ng'habi KR, Lee Y, Knols BG, Mwasheshi D, Lanzaro GC, Ferguson HM. Colonization of malaria vectors under semi-field conditions as a strategy for maintaining genetic and phenotypic similarity with wild populations. Malaria journal. 2015;14:10. Epub 2015/01/22. doi: 10.1186/s12936-014-0523-0. PubMed PMID: 25604997; PubMed Central PMCID: PMCPMC4340333.

57. Allgood DW, Yee DA. Influence of resource levels, organic compounds and laboratory colonization on interspecific competition between the Asian tiger mosquito *Aedes albopictus* (Stegomyia albopicta) and the southern house mosquito *Culex quinquefasciatus*. Medical and veterinary entomology. 2014;28(3):273–86. doi: 10.1111/mve.12047. PubMed PMID: 24444185; PubMed Central PMCID: PMC4105337.

58. Ochieng'-Odero JPR. Does laboratory adaptation occur in insect rearing systems, or is it a case of selection, acclimatization and domestication? Insect Science Applications. 1994;15(1):1–7.

59. Marchand R. A new cage for observing mating behavior of wild Anopheles gambiae in the laboratory. Journal of the American Mosquito Control Association. 1985;1(2):234.

60. Lardeux F, Quispe V, Tejerina R, Rodriguez R, Torrez L, Bouchite B, et al. Laboratory colonization of Anopheles pseudopunctipennis (Diptera: Culicidae) without forced mating. C R Biol. 2007;330(8):571–5. Epub 2007/07/20. doi: 10.1016/j.crvi.2007.04.002. PubMed PMID: 17637437.

61. McDaniel IN, Horsfall W. Induced copulation of aedine mosquitoes. Science (Washington). 1957;125.

62. Bryan JH, Southgate B. Studies of forced mating techniques on anopheline mosquitoes. Mosq News. 1978;38:338–42.

63. Ross PA, Axford JK, Richardson KM, Endersby-Harshman NM, Hoffmann AA. Maintaining *Aedes aegypti* mosquitoes infected with *Wolbachia*. Journal of Visualized Experiments. 2017;(126). doi: 10.3791/56124.

64. Jonas A, Taus T, Kosiol C, Schlotterer C, Futschik A. Estimating the effective population size from temporal allele frequency changes in experimental evolution. Genetics. 2016;204(2):723–35. Epub 2016/08/21. doi: 10.1534/genetics.116.191197. PubMed PMID: 27542959; PubMed Central PMCID: PMCPMC5068858.

65. Waples RS. A generalized approach for estimating effective population size from temporal changes in allele frequency. Genetics. 1989;121(2):379–91.

66. Jorde PE, Ryman N. Unbiased estimator for genetic drift and effective population size. Genetics. 2007;177(2):927–35. Epub 2007/08/28. doi: 10.1534/genetics.107.075481. PubMed PMID: 17720927; PubMed Central PMCID: PMCPMC2034655.

67. Economopoulos AP. Adaptation of the Mediterranean fruit-fly (Diptera: Tephritidae) to artificial rearing. Journal of Economic Entomology. 1992;85(3):753–8. PubMed PMID: WOS: A1992HW82700020.

68. Miyatake T. Difference in the larval and pupal periods between mass-reared and wild strains of the melon fly, Bactrocera-cucurbitae (Coquillett) (Diptera: Tephritidae). Applied Entomology and Zoology. 1993;28(4):577–81. PubMed PMID: WOS: A1993ML88900021.

69. Coon KL, Vogel KJ, Brown MR, Strand MR. Mosquitoes rely on their gut microbiota for development. Molecular ecology. 2014;23(11):2727–39. doi: 10.1111/mec.12771. PubMed PMID: 24766707; PubMed Central PMCID: PMCPMC4083365.

70. Coon KL, Brown MR, Strand MR. Gut bacteria differentially affect egg production in the anautogenous mosquito *Aedes aegypti* and facultatively autogenous mosquito *Aedes atropalpus* (Diptera: Culicidae). Parasites & vectors. 2016;9(1):375. doi: 10.1186/s13071-016-1660-9. PubMed PMID: 27363842; PubMed Central PMCID: PMCPMC4929711.

71. Mwadondo EM, Ghilamicael A, Alakonya AE, Kasili RW. Midgut bacterial diversity analysis of laboratory reared and wild Anopheles gambiae and *Culex quinquefasciatus* mosquitoes in Kenya. African Journal of Microbiology Research. 2017;11(29):1171–83.

72. Dickson LB, Ghozlane A, Volant S, Bouchier C, Ma L, Vega-Rua A, et al. Diverse laboratory colonies of *Aedes aegypti* harbor the same adult midgut bacterial microbiome. BioRxiv. 2017:200659. doi: 10.1101/200659.

73. Yeap HL, Mee P, Walker T, Weeks AR, O'Neill SL, Johnson P, et al. Dynamics of the “popcorn” *Wolbachia* infection in outbred *Aedes aegypti* informs prospects for mosquito vector control. Genetics. 2011;187(2):583–95. PubMed PMID: 21135075; PubMed Central PMCID: PMC3030498.

74. Powell JR, Evans BR. How much does inbreeding reduce heterozygosity? Empirical results from *Aedes aegypti*. The American journal of tropical medicine and hygiene. 2016;96(1):157–8. doi: 10.4269/ajtmh.16-0693. PubMed PMID: 27799643.

75. Koenraadt CJ, Kormaksson M, Harrington LC. Effects of inbreeding and genetic modification on *Aedes aegypti* larval competition and adult energy reserves. Parasites & vectors. 2010;3:92. doi: 10.1186/1756-3305-3-92. PubMed PMID: 20925917; PubMed Central PMCID: PMC2967506.

76. Armbruster P, Hutchinson RA, Linvell T. Equivalent inbreeding depression under laboratory and field conditions in a tree-hole-breeding mosquito. Proceedings Biological sciences / The Royal Society. 2000;267(1456):1939–45. Epub 2000/11/15. doi: 10.1098/rspb.2000.1233. PubMed PMID: 11075705; PubMed Central PMCID: PMCPMC1690768.

77. O'Donnell D, Armbruster P. Inbreeding depression affects life-history traits but not infection by *Plasmodium gallinaceum* in the Asian tiger mosquito, *Aedes albopictus*. Infection, genetics and evolution: journal of molecular epidemiology and evolutionary genetics in infectious diseases. 2010;10(5):669–77. doi: 10.1016/j.meegid.2010.03.011. PubMed PMID: 20359551.

78. Phuc HK, Andreasen MH, Burton RS, Vass C, Epton MJ, Pape G, et al. Late-acting dominant lethal genetic systems and mosquito control. BMC biology. 2007;5:11. doi: 10.1186/1741-007-5-11. PubMed PMID: 17374148; PubMed Central PMCID: PMC1865532.

79. Catteruccia F, Godfray HCJ, Crisanti A. Impact of genetic manipulation on the fitness of Anopheles stephensi mosquitoes. Science. 2003;299(5610):1225–7.

80. Shetty V, Shetty NJ, Harini BP, Ananthanarayana SR, Jha SK, Chaubey RC. Effect of gamma radiation on life history traits of *Aedes aegypti* (L.). Parasite Epidemiology and Control. 2016;1(2):26–35. doi: 10.1016/j.parepi.2016.02.007.

81. Endersby N, Hoffmann A, White V, Ritchie S, Johnson P, Weeks A. Changes in the genetic structure of *Aedes aegypti* (Diptera: Culicidae) populations in Queensland, Australia, across two seasons: implications for potential mosquito releases. Journal of Medical Entomology. 2011;48(5):999–1007.

82. Saarman NP, Gloria-Soria A, Anderson EC, Evans BR, Pless E, Cosme LV, et al. Effective population sizes of a major vector of human diseases, *Aedes aegypti*. Evolutionary Applications. 2017. doi: 10.1111/eva.12508.

83. Schou MF, Loeschcke V, Bechsgaard J, Schlotterer C, Kristensen TN. Unexpected high genetic diversity in small populations suggests maintenance by associative overdominance. Molecular ecology. 2017. Epub 2017/07/27. doi: 10.1111/mec.14262. PubMed PMID: 28746770.

84. Nunney L. Measuring the ratio of effective population size to adult numbers using genetic and ecological data. Evolution; international journal of organic evolution. 1995;49(2):389–92.

85. Hedrick P. Large variance in reproductive success and the Ne/N ratio. Evolution; international journal of organic evolution. 2005;59(7):1596–9.

86. Frankham R. Genetic rescue of small inbred populations: meta-analysis reveals large and consistent benefits of gene flow. Molecular ecology. 2015;24(11):2610–8. Epub 2015/03/06. doi: 10.1111/mec.13139. PubMed PMID: 25740414.

87. Menge DM, Guda T, Zhong D, Pai A, Zhou G, Beier JC, et al. Fitness consequences of Anopheles gambiae population hybridization. Malaria journal. 2005;4:44. Epub 2005/09/22. doi: 10.1186/1475-2875-4-44. PubMed PMID: 16174295; PubMed Central PMCID: PMCPMC1242248.

88. Ekechukwu NE, Baeshen R, Traore SF, Coulibaly M, Diabate A, Catteruccia F, et al. Heterosis increases fertility, fecundity, and survival of laboratory-produced F1 hybrid males of the malaria mosquito Anopheles coluzzii. G3 (Bethesda). 2015;5(12):2693–709. Epub 2015/10/27. doi: 10.1534/g3.115.021436. PubMed PMID: 26497140; PubMed Central PMCID: PMCPMC4683642.

89. Baeshen R, Ekechukwu NE, Toure M, Paton D, Coulibaly M, Traoré SF, et al. Differential effects of inbreeding and selection on male reproductive phenotype associated with the colonization and laboratory maintenance of Anopheles gambiae. Malaria journal. 2014;13(1):19.

90. Gilchrist AS, Meats AW. An evaluation of outcrossing to improve mass-reared strains of the Queensland fruit fly Bactrocera tryoni. International Journal of Tropical Insect Science. 2014;34(S1):S35–S44. doi: 10.1017/s1742758414000216.

91. Balestrino F, Puggioli A, Carrieri M, Bouyer J, Bellini R. Quality control methods for *Aedes albopictus* sterile male production. PLoS neglected tropical diseases. 2017;11(9):e0005881. Epub 2017/09/12. doi: 10.1371/journal.pntd.0005881. PubMed PMID: 28892483; PubMed Central PMCID: PMCPMC5608434.

92. Ritchie SA. Effect of some animal feeds and oviposition substrates on *Aedes* oviposition in ovitraps in Cairns, Australia. Journal of the American Mosquito Control Association-Mosquito News. 2001;17(3):206–8.

93. Rueda LM. Pictorial keys for the identification of mosquitoes (Diptera: Culicidae) associated with dengue virus transmission. Walter Reed Army Inst Of Research Washington Dc Department Of Entomology, 2004.

94. Ross PA, Endersby NM, Yeap HL, Hoffmann AA. Larval competition extends developmental time and decreases adult size of wMelPop *Wolbachia*-infected *Aedes aegypti*. The American journal of tropical medicine and hygiene. 2014;91(1):198–205. doi: 10.4269/ajtmh.13-0576. PubMed PMID: 24732463; PubMed Central PMCID: PMC4080562.

95. Livdahl TP, Sugihara G. Non-linear interactions of populations and the importance of estimating per capita rates of change. The Journal of animal ecology. 1984:573–80.

96. Axford JK, Ross PA, Yeap HL, Callahan AG, Hoffmann AA. Fitness of wAlbB *Wolbachia* infection in *Aedes aegypti*: parameter estimates in an outcrossed background and potential for population invasion. The American journal of tropical medicine and hygiene. 2016;94(3):507–16. doi: 10.4269/ajtmh.15-0608. PubMed PMID: 26711515.

97. Segoli M, Hoffmann AA, Lloyd J, Omodei GJ, Ritchie SA. The effect of virus-blocking *Wolbachia* on male competitiveness of the dengue vector mosquito, *Aedes aegypti*. PLoS neglected tropical diseases. 2014;8(12):e3294.

98. Schneider CA, Rasband WS, Eliceiri KW. NIH Image to ImageJ: 25 years of image analysis. Nature methods. 2012;9(7):671–5.

99. Rašić G, Filipović I, Weeks AR, Hoffmann AA. Genome-wide SNPs lead to strong signals of geographic structure and relatedness patterns in the major arbovirus vector, *Aedes aegypti*. BMC genomics. 2014;15(1):275.

100. Schmidt TL, Filipović I, Hoffmann AA, Rašić G. Fine-scale landscape genomics helps explain the slow spread of *Wolbachia* through the *Aedes aegypti* population in Cairns, Australia. biorxiv. 2017.

101. Andrews S. FastQC: a quality control tool for high throughput sequence data. 2010.

102. Catchen J, Hohenlohe PA, Bassham S, Amores A, Cresko WA. Stacks: an analysis tool set for population genomics. Molecular ecology. 2013;22(11):3124–40. Epub 2013/05/25. doi: 10.1111/mec.12354. PubMed PMID: 23701397; PubMed Central PMCID: PMCPMC3936987.

103. Dudchenko O, Batra SS, Omer AD, Nyquist SK, Hoeger M, Durand NC, et al. De novo assembly of the *Aedes aegypti* genome using Hi-C yields chromosome-length scaffolds. Science. 2017;356(6333):92–5.

104. Langmead B, Salzberg SL. Fast gapped-read alignment with Bowtie 2. Nat Methods. 2012;9(4):357–9. Epub 2012/03/06. doi: 10.1038/nmeth.1923. PubMed PMID: 22388286; PubMed Central PMCID: PMCPMC3322381.

105. Li H, Handsaker B, Wysoker A, Fennell T, Ruan J, Homer N, et al. The sequence alignment/map format and SAMtools. Bioinformatics. 2009;25(16):2078–9.

106. Kofler R, Pandey RV, Schlotterer C. PoPoolation2: identifying differentiation between populations using sequencing of pooled DNA samples (Pool-Seq). Bioinformatics. 2011;27(24):3435–6. doi: 10.1093/bioinformatics/btr589. PubMed PMID: 22025480; PubMed Central PMCID: PMCPMC3232374.

